# Inferring Thalamocortical Monosynaptic Connectivity In-Vivo

**DOI:** 10.1101/2020.10.09.333930

**Authors:** Yi Juin Liew, Aurélie Pala, Clarissa J Whitmire, William A Stoy, Craig R Forest, Garrett B Stanley

**Affiliations:** Wallace H Coulter Department of Biomedical Engineering, Georgia Institute of Technology and Emory University, Atlanta, GA, USA; George W Woodruff School of Mechanical Engineering, Georgia Institute of Technology, Atlanta, GA, USA

**Author notes:** Correspondence: Garrett B Stanley Coulter Department of Biomedical Engineering Georgia Institute of Technology & Emory University 313 Ferst Dr Atlanta GA 30332-0535, USA. **Disclosures:** The authors declare no competing financial interests.

**Keywords:** thalamocortical, cross correlation, signal detection, causality, inference

## Abstract

As the tools to simultaneously record electrophysiological signals from large numbers of neurons within and across brain regions become increasingly available, this opens up for the first time the possibility of establishing the details of causal relationships between monosynaptically connected neurons and the patterns of neural activation that underlie perception and behavior. Although recorded activity across synaptically connected neurons has served as the cornerstone for much of what we know about synaptic transmission and plasticity, this has largely been relegated to *ex-vivo* preparations that enable precise targeting under relatively well-controlled conditions. Analogous studies *in-vivo*, where image-guided targeting is often not yet possible, rely on indirect, data-driven measures, and as a result such studies have been sparse and the dependence upon important experimental parameters has not been well studied. Here, using in-vivo extracellular single unit recordings in the topographically aligned rodent thalamocortical pathway, we sought to establish a general experimental and computational framework for inferring synaptic connectivity. Specifically, attacking this problem within a statistical signal-detection framework utilizing experimentally recorded data in the ventral-posterior medial (VPm) region of the thalamus and the homologous region in layer 4 of primary somatosensory cortex (S1) revealed a trade-off between network activity levels needed for the data-driven inference and synchronization of nearby neurons within the population that result in masking of synaptic relationships. Taken together, we provide a framework for establishing connectivity in multi-site, multi-electrode recordings based on statistical inference, setting the stage for large-scale assessment of synaptic connectivity within and across brain structures.

**New & Noteworthy:** Despite the fact that all brain function relies on the long-range transfer of information across different regions, the tools enabling us to measure connectivity across brain structures are lacking. Here, we provide a statistical framework for identifying and assessing potential monosynaptic connectivity across neuronal circuits from population spiking activity that generalizes to large-scale recording technologies that will help us to better understand the signaling within networks that underlies perception and behavior.

## Introduction

Every aspect of brain function, from sensory and motor processing, to memory and cognition, involves complex circuitry and communication across many different brain areas. Despite this fact, what we know about brain function has been derived largely from electrophysiological recordings targeted at single regions, and upon gross anatomical connection patterns across brain regions without specific precise knowledge of synaptic connectivity. Ultimately, understanding the causal interaction of neuronal dynamics that underlie perception and behavior requires ground truth evidence for synaptic connectivity that necessitates intracellular access to both pre- and post-synaptic neurons. Despite exciting advancement of a range of recording technologies such as simultaneous multi-neuron intracellular recordings in-vivo (Jouhanneau and Poulet 2019; Kodandaramaiah et al. 2018) and deep structure targeted patching (Stoy et al. 2017), targeting connections in-vivo using intracellular approaches remains a labor-intensive endeavour, is ultimately limited to a very small number of neurons, and does not scale to the circuit level. Extracellular recordings offer solutions to some of these issues, and thus offer promise in this direction.

The field of neuroscience is now at a time where large-scale recordings of neuronal populations at cellular resolution are possible across brain structures, due to the development of multi-electrode recording technologies such as ‘Neuronexus’ (Berényi et al. 2014; Buzsáki et al. 2015) (NeuroNexus Inc.), ‘Neuropixels’ (Jun et al. 2017), ‘NeuroSeeker’ (Fiáth et al. 2018; Raducanu et al. 2016)), and 3D silicon probes (Rios et al. 2016). These technologies provide access to surveying network level information flow, driving a growing need for a rigorous experimental and analytic framework to identify functional relationships across brain structures. In previous studies, in-vivo approaches to establishing synaptic connectivity across recorded pairs of neurons have been developed based on analytic methods applied to recorded spiking data (Nowak and J 2000; Perkel et al. 1967), historically conducted in larger animals such as cats, rabbits, and rats (Reid and Alonso 1995; Swadlow and Gusev 2001; Wang et al. 2010) and only more recently in mouse (Lien and Scanziani 2018). Approaches based on spike correlations are the most common for quantifying functional interactions and inferring monosynaptic connections in-vivo. These approaches have been utilized in paired recording studies that involved measuring correlation in the spiking activity between neurons (Barthó et al. 2004; Csicsvari et al. 1998; English et al. 2017; Fujisawa et al. 2008; Reid and Alonso 1995; Swadlow and Gusev 2001; Wang et al. 2010) or between the spiking activity of a pre-synaptic neuron and the subthreshold membrane potential of a putative post-synaptic target (Bruno and Sakmann 2006; Matsumura et al. 1996) (London et al. 2010; Sedigh-Sarvestani et al. 2017; Yu and Ferster 2013). Correlational approaches are statistical in nature and thus have been anecdotally reported to be strongly dependent upon data length and are very sensitive to a range of possible confounds, but a comprehensive understanding of these relationships and a unified approach are both lacking.

Here, using the thalamocortical circuit in rodents as a model system, we establish methodological strategies for extracellular, topographically aligned in-vivo single-unit recordings in thalamus and cortex for both rat and mouse. This analytic framework for inferring connectivity is based on signal-detection theory, and directly addresses issues of data length dependence and confounds produced by population synchrony. Specifically, we outline a work-flow of topographic mapping and histological validation, followed by a statistical approach for signal-detection based classification of putative connected and non-connected pairs with an assessment of classification confidence that is scalable to the large-scale recording approaches that are emerging in the field. We found that while the amount of spiking was a strong determinant of the accuracy of the inference of connectivity, there was an important tradeoff between the activity in the network and the underlying population synchrony that regulates the likelihood of both discovering a synaptic connection that is present and correctly classifying unconnected pairs as such. Taken together, we provide a data-driven framework for inferring connectivity and the corresponding statistical confidence that generalizes to large-scale recordings across brain structures.

## METHODS

### Animals

Twelve female adult Sprague Dawley rats (no age restriction, 225-330g) and four adult male C57BL/6 mice (8-16 weeks, 25-35g) were used in all the experiments. All procedures were approved by the Georgia Institute of Technology Institutional Animal Care and Use Committee and followed guidelines established by the National Institutes of Health. Note that there were some major differences in animal preparation, surgery and craniotomies as well as electrophysiology for rat and mouse, as described below. However, procedures such as whisker stimulation, post-mortem histology, and analytical methods used for analysis were largely identical.

### Animal preparation, surgery and craniotomies

#### Rat

Animals were first sedated with 5% vaporized isoflurane and maintained at 3% vaporized isoflurane when transitioned to fentanyl-cocktail anesthesia via tail vein injection. Fentanyl cocktail anesthesia (fentanyl [5 μg/kg], midazolam [2 mg/kg], and dexmedetomidine [150 mg/kg])) was administered continuously via a drug pump at an initial rate of 4.5 uL/min (Ahissar et al. 2000; Whitmire et al. 2016). During the transition, anesthesia level was monitored closely by measurement of heart rate and respiratory rate. Body temperature was maintained at 37 °C by a servo-controlled heating pad. Under the effect of both anesthetic agents, the animal’s heart rate tended to decrease gradually over several minutes. The isoflurane level was then titrated down in 0.5% decrements during the transition to the fentanyl cocktail. Upon successful transition, which ranged from 5-15 min depending on the animal, the heart rate was targeted for approximately 240 -270 bpm. After the effects of fentanyl cocktail stabilized (in terms of heart rate) and the animals showed no toe pinch response, the animals were then fixed in the stereotaxic device by securing the head of the animal in place with ear bars on a floating table in an electromagnetically shielded surgery suite. The position of the ear bars was verified with uniform eye levels and usually with eardrum penetration on both sides of the head. Eye ointment (Puralube Vet Ointment) was applied to prevent the animal’s eyes from dehydration. An incision was made along the midline of the skull, and skin was removed for visibility of bregma and lambda (Paxinos and Watson 2007). Connective tissue and muscle that were close to the ridge of the skull were detached and removed to expose the skull surface that was directly above the barrel cortex. To ensure that the skull remained level throughout the recording, the height of the skull surface was measured at bregma and lambda, and the difference was minimized (< 200 um) by adjusting the angle of the head with the nose cone position.

We made two craniotomies on the animal’s left hemisphere above the ventro-posteromedial nucleus (VPm) and the S1 barrel cortex based on stereotaxic coordinates. For the cortical craniotomy, which usually extended over the ridge of the skull, the stereotaxic device was rotated to an angle (∼40° relative to vertical) for better visualization and drilling. Both craniotomies were drilled slowly until the skull piece appeared to be floating and mobile. We irrigated the skull surface periodically with Ringer’s solution when drilling to remove debris and prevent overheating from drilling. Before removing the skull piece to expose the cortex, a small dental cement reservoir was carefully built around the craniotomies to hold the Ringer’s solution for irrigation and to keep the recording site continually moist. The lateral side of the wall was built thicker and higher due to the curvature of the skull toward the ridge. To provide strong adherence to the skull, cyanoacrylate containing adhesive (Krazy Glue, Elmer’s Products Inc.) was carefully applied around the external edges of the reservoir, leaving the bregma and major skull sutures visible at all times. For grounding purposes, another hole with ∼0.5 mm diameter was drilled (Henry Schein, Carbide Burr HP 2) on the right hemisphere of the skull and fastened with a skull screw and a metal wire. The skull pieces at both recording sites were then carefully removed with forceps (FST Dumont #5/45). To minimize brain swelling, the dura was left intact on both recording surfaces, and warm Ringer’s solution was repeatedly added and absorbed with cotton tips until the blood was cleared. An absorbent gelatin compressed sponge (Gelfoam, Pfizer Inc.) was sometimes used to clear the blood.

#### Mouse

Head-plate implantation and intrinsic imaging procedure were usually performed at least three days to one week before acute experiment. A lightweight custom metal (titanium or stainless steel) head-plate was implanted on the skull of the mouse for head fixation and improved stability during recording, in accordance with a previously described protocol (Borden et al. 2017). During this survival surgical preparation, the animal was sedated with 5% vaporized isoflurane and anesthesia was maintained with 2-3% isoflurane for the head-plate procedure. We administered opioid and non-steroidal anti-inflammatory analgesic (SR-Buprenorphine 0.8 - 1 mg/kg, SC, pre-operatively, and Ketoprofen 5-10 mg/kg, IP, post-operatively) and covered the animal’s eyes with ophthalmic eye ointment (Puralube Vet Ointment). Body temperature was monitored and maintained at 37 °C. After sterilizing the skin above the skull and applying topical anesthetic (lidocaine 2%, 0.05 ml, max 7mg/kg, SC), we removed the skin and scraped off periosteum and any conjunctive tissue off the skull. We gently separated tissue and muscles close to the lateral edge of the skull using a scalpel blade (Glass Van & Technocut, size no. 15), leaving sufficient room for head-plate attachment and away from targeted recording areas. To ensure that the skull surface remained level during head fixation, we adjusted animal’s head position to minimize the relative height difference (< 150 um) between the skull surface at the bregma and lambda landmarks (Franklin and Paxinos 2008). We found this step to be critical especially for VPm targeting. We secured the metal headplate over the intact skull with C&B-Metabond (Parkell Prod Inc) and skin adhesive (Loctite 401, Henkel). The Metabond dental acrylic was chilled using ice, slowly applied to the skull surface, and allowed to cure for 5-10 minutes before we covered the rest of the attachment site and edges of the skin incision with skin adhesive (Loctite 401) to ensure that the skin edges were securely adhered to the skull. The final head-plate and dental acrylic structure created a recording well for holding Ringer solution (in mM: 135 NaCl, 5 KCl, 5 HEPES, 1 MgCl_2_-6H_2_O, 1.8 CaCl_2_-2H_2_Ol pH 7.3) for future imaging and electrophysiological recording sessions. For grounding purposes, another hole with ∼0.5 mm diameter was drilled (Henry Schein, Carbide Burr HP 2) on the right hemisphere of the skull and fastened with a skull screw (Miniature Self-Tapping Screws, J. I. Morrisco). A metal wire was connected to the skull screw on the day of recording to serve as animal ground. We applied a thin layer of transparent glue (Loctite 401, Henkel) over the left hemisphere to protect the skull and covered the exposed skull with a silicone elastomer (Kwik-Cast, World Precision Instruments) if there were no other additional procedures after head-plate implantation.

##### Intrinsic Optical Imaging in the Mouse S1 Barrel Cortex

We performed intrinsic signal optical imaging of whisker-evoked responses in mouse primary somatosensory cortex under 1-1.2% isoflurane anesthesia to functionally identify individual barrel columns (Lefort et al. 2009; Pala and Petersen 2015). All whiskers except the whiskers of interest (A2, B1, C2, D1, D3 and E2) were trimmed. We thinned the skull using a dental drill (0.1 mm diameter bit, Komet, USA) until blood vessels became visible. We applied warm Ringer’s solution on top of the skull and covered it with a glass coverslip (thickness 0.13-0.17mm, Fisherbrand). We captured a reference image of the blood vasculature under green illumination (530nm) (M530F1 LED, Thorlabs). We delivered repetitive whisker stimuli (10 Hz, sawtooth pulses, 1000 deg/s) to individual whiskers using a galvanometer system (Cambridge Technologies) while performing imaging on barrel cortex under red illumination (625 nm) (M625F1 LED, Thorlabs) – see Whisker Stimulation section below. Images were acquired using a CCD Camera (MiCam02HR, SciMedia) at 10 Hz, with a field of view of approximately 4 x 2.5 mm, corresponding to 384 x 256 pixels (resolution: 100pixel/mm). Each trial lasted 10s, with 4s of baseline followed by 4s of whisker stimulation and 2s without stimulation. The inter-trial interval was 30s. Whisker-evoked responses were averaged over 10-20 trials. Intrinsic signals were measured as the relative change in reflectance by taking the overall mean reflectance during whisker stimulation (R_stim_) and subtracted the mean baseline reflectance (R_stim_ - R_baseline_). Acquisition and processing of the images were implemented using “BV_Ana” software (MiCam02HR, SciMedia, Ltd) and Matlab (MathWorks, Natick, 2015). We recorded intrinsic signals from at least three barrels and estimated location of the unmapped barrels by overlaying a template barrel map reconstructed from histology. At the end of the imaging session, we applied transparent glue (Loctite 401) directly on the S1 site to protect the thinned skull from contamination and infection and sealed the exposed skull with a silicone elastomer (Kwik-Cast, World Precision Instruments).

On the day of recording, the head of the mouse was held in place with a head-post clamp and animal should remain in the flat skull position. We covered the eyes with ophthalmic eye ointment (Puralube Vet Ointment) to prevent dehydration. The animal was then anesthetized using 2-3% isoflurane. Before performing the craniotomies, we thinned the skull above VPm and S1 layer-by-layer with a dental drill (0.1 mm diameter bit, Komet, USA) and irrigated the skull surface with Ringer’s solution frequently to remove debris and prevent overheating. When the skull was thin enough to easily puncture, we made two small craniotomies above VPm (approximately 1 mm diameter; centered at 1.8 mm caudal and 1.8 mm lateral to bregma) and S1 (approximately 0.5 mm diameter, using the intrinsic imaging signal and the blood vasculature as a landmark) by penetrating the thinned skull using an insulin syringe needle tip (at about 40-50 degree angle) to make small holes outlining the edges of the craniotomies until the circular skull piece was loosely attached. The less perforated region of the circular skull piece was used as a pivot, and we levered the skull pieces off from the more perforated edge using fine-tip forceps (FST Dumont #5SF) to expose the brain surface.

### Electrophysiology

#### Neuronal recordings

Tungsten microelectrodes (FHC, impedance: 2-5 MΩ, 75 µm in diameter) were used in thalamus and cortex to isolate single units that were responding to a single primary whisker on the contralateral side of the face. Multielectrode silicon probes with 32 recording sites (A1×32-Poly3-5mm-25s-177 or A1×32-Poly2-10mm-50s-177, NeuroNexus) were also independently lowered into the thalamus and cortex. To improve signal quality, we electroplated the silicon probes with the polymer PEDOT:PSS (Baião 2014) with a nanoZ device (Multi Channel Systems, Germany). The impedances of the contact sites were measured prior to each recording and ranged from 0.3-0.8 MOhms. After each use, the contact sites were soaked in 20% Contrad (labware detergent) overnight for cleaning and the impedances usually returned to the initial values (0.3-0.8 MOhms). The impedances for the silicon probes were always measured prior to a recording session and contact sites that were defective usually showed large fluctuations in raw voltage traces and signals detected at these sites were excluded for analysis prior to spike-sorting. Note that tungsten microelectrodes were used in rat studies and probe recordings were used in mouse studies. Data were collected using a 64-channel electrophysiological data-acquisition system (TDT Model, etc. RZ2 Bioprocessor). Neuronal signals were amplified, and bandpass filtered (500Hz-5kHz) and digitized at ∼25kHz per channel. Stimulus waveform and other continuous data were digitized at 10kHz per channel. Simultaneously, the local field potential (LFP) signals were obtained by using a 0.5-200Hz bandpass filter. The LFPs were used to identify primary whisker response (see Whisker Stimulation).

#### Rat

*VPm recordings.* We targeted the VPm region of the thalamus by advancing a single tungsten microelectrode (FHC, 2 MOhm impedance) perpendicular to the pial surface into the thalamic craniotomy centered at 3.0 mm caudal and 3.0 mm lateral to bregma. We quickly advanced (∼50um/s) to the depth of 4000 µm and slowed down to the speed of ∼3 µm/s while searching for responsive cells by manually deflecting the whisker on the contralateral side of the vibrissal pad. Whisker responsive cells were typically located at a depth of 4800-5300 µm, measured using a precision micromanipulator (Luigs & Neumann, Germany). *S1 recordings*. At a position 5.8 mm lateral and 2.5 mm caudal to bregma, we inserted a single tungsten microelectrode at an angle of 40-45° (relative to the vertical axis of electrode holder). We positioned the tip of the electrode gently touching the dura and advanced slowly to create an opening in the membrane. The microelectrode was then slowly advanced into cortical tissue at a speed of ∼3 µm/s (measured at 1µm resolution). All cortical units were recorded at stereotaxic depths of 700-1000µm, corresponding to layer IV of rat barrel cortex based on literature (Bruno and Sakmann 2006; Constantinople and Bruno 2013).

#### Mouse

*VPm recordings.* For first penetration, electrode was typically lowered at 1.8 mm caudal and 1.8 mm lateral to bregma. If we could not functionally locate VPm using this method, we used the relative distance from the location of barrel cortex (usually measured 1.0-1.2mm from the B1 or Beta barrel column to find whisker responsive thalamic regions). For probe recordings, we searched for whisker responsive cells by manual whisker deflection using the deepest channel as reference. We typically found whisker responsive cells were at a depth between 2800um-3200 um. *S1 recordings.* Based on intrinsic imaging signal, a small craniotomy (diameter ∼0.5 mm) was created at the desired location on the barrel cortex that was distant from blood vessels. We positioned the electrode at a 35 ° angle from the vertical axis (parallel to the barrel column) and recorded from cortical neurons at a stereotaxic depth of 300-500um from the cortical surface.

### Whisker Stimulation

To identify the single whisker of interest, we first manually deflected all whiskers on the vibrissal pad while monitoring the extracellular signal. Once units were isolated by moving the electrode as close as possible to the responsive cells that primarily responded to single whisker, we delivered controlled, single whisker stimulation in the rostral-caudal plane using a computer-controlled actuator, galvo-motor (galvanometer optical scanner model 6210H, Cambridge Technology). Whiskers were trimmed ∼12 mm from the face. Primary whisker (PW) was defined as whisker deflection that evoked the maximal neuronal response with shortest latency (see Transient Stimulus). The primary whisker was fed into the insertion hole at the end of an extension tube (inner diameter: ∼200-300 µm um, length: 15mm) that was connected to the rotor of the galvanometer stimulator. The stimulus probe was positioned at 10 mm from the vibrissal pad. The range of motion of the galvanometer was ±20°, with a bandwidth of 200Hz. The galvanometer system was controlled using a custom developed hardware/software system (Matlab Real-Time Simulink System, MathWorks). Whisker evoked responses were measured using raster plots and peri-stimulus time histograms (PSTH, 1-ms bin resolution) across trials.

#### Transient stimulus

For mapping whisker receptive field at the recording sites, we presented high velocity (600 or 1000 °/s) rostral-caudal whisker deflection to evoke reliable whisker responses. Trains of solo pulses, followed by a separate, pulsatile adapting stimulus at 8-10 Hz (Borden et al. 2017; Whitmire et al. 2016) were delivered for 60-100 times. To measure whisker response to different stimulus strength, pulses with angular deflection velocities of 50, 125, 300, 600, 900, 1200 deg/s were presented randomly. Rostral-caudal pulse deflections were either a Gaussian-shaped deflection waveform or a simple exponential sawtooth (rise and fall time = 8 ms). We quantified whisker evoked activity 30 ms after stimulus onset and then further characterized onset latencies of first spike within the response window in response to non-adapting solo pulses and adapting pulsatile stimuli over 60-100 trials. Mean first spike latencies, defined as the average time delay between stimulus onset and the first spike in the response window after stimulus presentation. This metric was used for all thalamic and cortical units. For 32-channel probe recording in thalamus, we quantified the LFP response in a 30 ms post-stimulus window for each channel to identify principal whiskers (PW). Well-isolated single units that showed whisker responsiveness was verified by calculating LFP peak amplitude and peak latency to different whisker stimulation.

A neuronal pair was verified to be topographical aligned when they (1) shared maximum response (SUs max FR and LFP amplitude) to the same primary whisker under punctate stimulation and (2) latency difference of whisker response within 1-5 ms delay (LFP peak latency and SU mean FSL, or SU mean peak latency).

#### Weak sinusoidal stimulus

For monosynaptic connection quantification, we probed the thalamocortical circuit using a weak, desynchronizing sinusoidal whisker stimulus to elevate baseline firing rates in recorded units. We delivered 4°, 2-4Hz weak sinusoidal deflections (Bruno and Sakmann 2006; Bruno and Simons 2002; Wang et al. 2010) for approximately 200-500 trials to obtain at least 2000 spikes for the cross-correlation analysis. Occasionally, we eliminated the first 0.5-1 s of the trials due to high firing rate at the onset of stimulus presentation.

### Post-mortem Histology

In order to verify recording sites and the angle of penetration that was optimal for locating VPm and S1, we perfused a small subset of animals after the paired recording experiment. To label each electrode recording track, we slowly retracted the electrode along its axis of entry at the conclusion of recording and applied a few drops of DiI (2.5mg/mL, in ethanol). We then reinserted the electrode into the same penetration site and back along the same axis and left the electrode in the brain for at least 15 minutes. Following the end of the experiment, we euthanized the animal with an overdose of sodium pentobarbital (euthasol, 0.5 mL at 390mg/mL for rat, 0.1 mL at 390 mg/mL), performed a transcardial perfusion, extracted the brain, and fixed the brain in 4% paraformaldehyde (PFA, Electron Microscopy Sciences) overnight. We sliced the brain in 100 um coronal sections and performed cytochrome oxidase staining to reveal VPm (barreloid) and S1 (barrel). As an additional verification of recording site, we identified the overlap between the DiI stained recording track and the CO-stained regions.

### Analytical Methods

#### Extracellular spike sorting

In assessing potential synaptic connectivity using cross-correlation analysis, the clear isolation of single unit activity from extracellularly recorded voltage signals is particular critical. In all of our paired recording experiments, we performed extracellular single-unit recordings using either Tungsten microelectrodes or 32-channel NeuroNexus probes. Although similar in approach, the sorting of recorded data into clusters were implemented using different software. For the single micro-electrodes, we performed spike sorting offline using the Plexon Offline Sorter (Plexon Inc, Dallas TX, USA), while for the 32-channel probe recordings we utilized KiloSort2 software package (https://github.com/MouseLand/Kilosort2) (Harris et al. 2000). For high density probe recordings, there was an additional manual curation step using phy (https://github.com/cortex-lab/phy) where we refined the output of automatic algorithm and determined if merging or splitting of specific clusters were necessary based on refractory violation, waveform shape as well as cross-correlogram between clusters. For both Tungsten microelectrode and high-density probe recordings, we classified the clusters as single-or multi-units based on the signal-to-noise ratio and inter-spike interval (ISI) distribution. We only included well-isolated clusters for further analysis. The selection of well-isolated clusters, or single units, was based on two criteria: (1) high signal-to-noise ratio (SNR) of the spike waveform: Peak-to-peak amplitude (Vpp) of spike waveforms greater than three standard deviation (SD) of the waveform. (2) Had a clear refractory period(Buzsáki 2004; Fiáth et al. 2019). We defined a single-unit as well-isolated if it had SNR greater than 3, ISI violation less than 1% for cortical unit and 2% for thalamic unit and spike waveform with a peak-to-peak amplitude (Vpp) greater than 60 µV. Given that the spikes of the same neuron were usually detected on multiple sites of a 32-channel probe, the peak-to-peak amplitude and standard deviation of the waveform were computed using channel that had the largest amplitude of spike waveform.

#### VPm unit verification

It is important to distinguish different thalamic nuclei from one another, and particularly so in the mouse, where the regions are very small and very close together. After recording, we used a combination of measures to classify thalamic units as VPm: (1) Average first-spike latency to punctate (non-adapting) stimuli, quantifying the average time between stimulus onset and the first spike fired during the neural response window (30 ms after stimulus onset) (Storchi et al. 2012). (2) Shift in average first-spike latency response to adapting stimuli, defined as the time difference between average response to the first adapting stimulus and last adapting stimulus (Masri et al. 2008). (3) Response reliability, defined as the percentage of trials where a response was detected within 20ms to repetitive stimulation (8-10Hz) (Mainen and Sejnowski 1995). We excluded thalamic units with average first-spike latency of more than 12 ms (see Supplementary Figure 2A), latency shift of more than 20ms, or response reliability of less than 20%.

#### Cortical unit classification

It was known that S1 layer IV is the major thalamocortical recipient from VPm projections and there exist some heterogeneity in terms of cortical cell types in Layer IV (Bruno and Simons 2002). Here, we classified cortical units into fast-spiking unit (FSU) and regular-spiking unit (RSU) by using waveform parameter. We quantified the time interval between trough to peak (t_t2p_) of the spike waveform (Barthó et al. 2004).

#### Cross-correlation analysis

All analysis was performed in a trial-by-trial basis. Given a spike in a ‘reference’ neuron, we computed the relative times of the spikes from a ‘target’ neuron that occurred within a 25-millisecond window before and after each reference spike. Cross-correlograms were constructed using a 0.5 milliseconds bin. Traditionally, monosynaptic interactions were known to produce short latency peaks in the cross-correlograms. The latency of the monosynaptic peak within cross-correlograms was typically reported to center around 2.5 ms, estimated from thalamocortical EPSPs and spiking activity (Alonso and Martinez 1998; Alonso et al. 2001; Bruno and Sakmann 2006; Bruno and Simons 2002; Reid and Alonso 1995; Sedigh-Sarvestani et al. 2017; Swadlow et al. 1978; Swadlow 1989; 2003; Swadlow and Gusev 2001). To ensure our analysis captures correlation from monosynaptic delays only, we allowed for 1.5 ms jitter on each side, setting the lower bound to be 1 ms and the upper bound to be 4 ms. This eliminated peaks that could arise in the 0-1 ms bin due to a shared common input, as well as disynaptic EPSPs that could have latencies longer than approximately 5 ms (Gil and Amitai 1996). Here, for the monosynaptic connectivity inference, we used thalamic spiking as the reference and cortical spiking as the target and examined correlated firing within a lag window of 1-4 ms. The results here were relatively invariant to the specific choice over a range of lag windows, but we found the 1-4ms window to be the most conservative (i.e. smaller window than 1-5 ms but resulted in same number of pairs classified as “connected”, see Supplementary Figure 3). For the spontaneous condition, we segmented the spike train data into 5s-trials (matching duration of stimulus-based trials) and performed cross-correlation analysis in a trial-by-trial manner. To measure the level of presynaptic synchronization in thalamus, we performed a cross-correlation analysis and computed synchrony strength using a central area under the cross-correlogram for all thalamic pairs that responded to the same principal whisker. The principal whisker was identified using the functional responses as measured by the local field potential (LFP) and in the single unit (SU) data. Thalamic synchrony was computed for each thalamic pair using the total number of spikes within a 15 ms (± 7.5 ms) window of the cross-correlogram (N_cc_), normalized by the mean number of spikes from each neuron (N_ref_, N_target_) (Alonso and Martinez 1998; Bruno and Sakmann 2006; Temereanca et al. 2008; Wang et al. 2010; Whitmire et al. 2016):

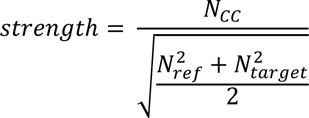

#### Probabilistic measure of connectivity inference

For each pair of thalamocortical neurons, we repeated the cross-correlational analysis with bootstrapping method. This was done by performing resampling on the dataset with replacement, with approximately 500-1000 iterations for each condition. By counting the number of iterations that fulfill the criteria for classifying the monosynaptic connection, this resulted in a p-value that can be attached to the binary classification of monosynaptic connection (see Results: Inferring connectivity in the context of a signal detection framework). We referred to this p-value as the probability of inferring monosynaptic connection, P(Inferring ‘connected’).

##### Data-length effect on the connectivity metric

Simulation for various data-lengths was implemented by using subsamples (in unit of trials) of VPm and S1 spikes for cross-correlation analysis. For each data-length condition, we computed the resampled peak height, h*. A bootstrapped estimator of bias was computed as the difference between the mean of the resampled peak height and the original metric, assuming a normal distribution:

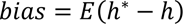

Where E(.) denotes statistical expectation. The variance was estimated as the square of the resampled peak height:

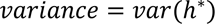

A significance level for the inference for a ‘putatively connected’ pair was computed and denoted as the probability of a hit when the peak height metric for the resampled data exceeded the criterion for the inference of a functional connection. The probability of a miss is complementary:

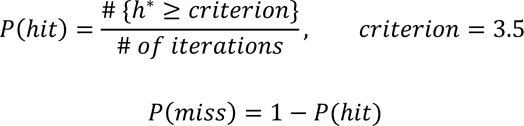

On the other hand, the significance level for a ‘not connected’ pair was computed and denoted as the probability of a correct reject when the peak height metric for the resampled data is less than the criterion.

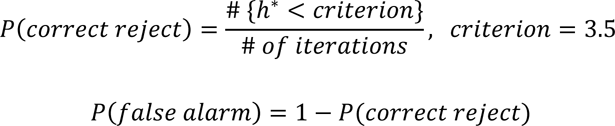

##### Thalamic synchrony effect on connectivity metric

Simulation for various thalamic synchrony level was implemented by manipulating both firing rate and spike timing of VPm and S1. Given that both firing rate and thalamic synchrony were affected with increasing stimulus strength, our simulation was conducted using data that was collected under spontaneous condition. For a connected pair, we set the ratio between VPm and S1 firing to be constant at each synchrony level for simplification. However, we systematically increased spiking activity (in number of spikes per trial) in VPm and S1 with a specific amount of jitter (zero-mean Gaussian noise of a specific standard deviation, sigma) as a function of thalamic synchrony. For a not-connected pair, the increase in thalamic synchrony and VPm and S1 firing were simulated by systematically adding spikes from another pair of neurons that is putatively connected. The assumption here was that a not-connected pair would become more identical to its neighboring neuron (that has a functional connection to the downstream S1 neuron) with increasing synchrony. This was repeated from 100-500 iterations for each synchrony level and the probability of satisfying Criterion 1 and 2 was computed.

### Results

Here, we present a comprehensive experimental and analytic framework for assessing synaptic connectivity using extra-cellular spiking activity from simultaneously recorded single units across brain structures. First, we provide a rigorous experimental protocol for performing simultaneous single-unit electrophysiological recording in topographically aligned regions in the thalamocortical circuit of the somatosensory pathway using a combination of stereotaxic targeting based on anatomical landmarks, sensory-evoked response properties, and post-hoc histological validation. We highlight some similarities and differences of the technical aspects of paired recordings performed in rats versus mice, two commonly used rodent species in mammalian electrophysiology. Next, we generate inferences regarding the synaptic connectivity of two single units based on spike correlation analysis across the presynaptic VPm and postsynaptic S1 spiking, leading to classification as either “connected” or “not connected”, using an approach to be explained in detail below. Figure 1A highlights the basic experimental setup and analysis utilized in this study, with electrophysiological recordings targeted to primary somatosensory “barrel” cortex (S1, red) and to ventro-posteromedial thalamus (VPm, blue), during controlled deflections of a single whisker on the contralateral side of the face with a computer-controlled actuator (see Methods). In the context of a signal detection framework, we then quantify the effects of experimental data-length and local synchrony of presynaptic neurons on monosynaptic connection inference, and importantly, expand this framework to attach statistical levels of confidence to the synaptic connectivity inferences.

**Figure 1:**
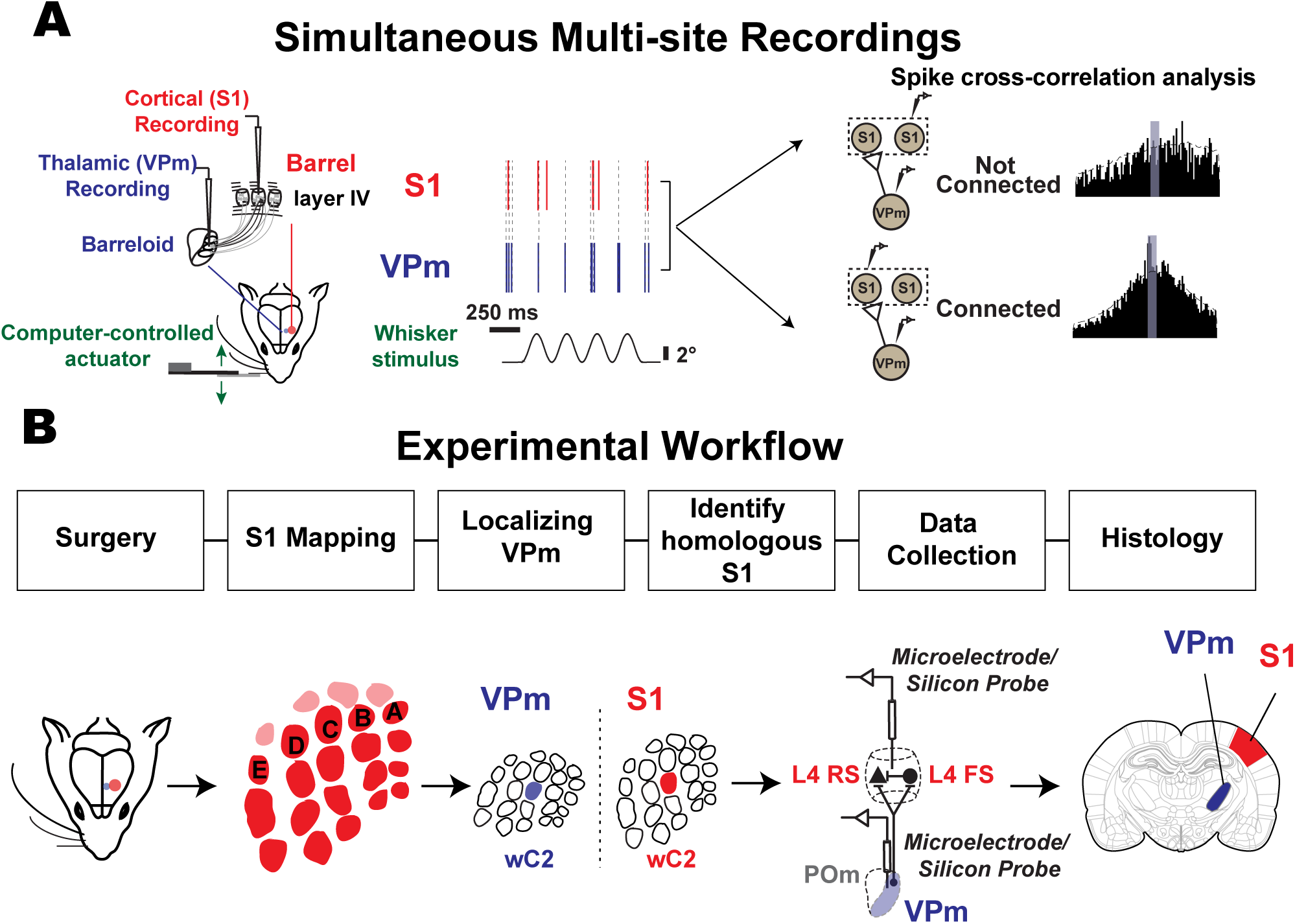
Experimental approach used to estimate monosynaptic connectivity between somatotopically - organized areas of the rodent somatosensory pathway. **A.** Simultaneous single-unit extracellular recordings were performed in the ventral posteromedial (VPm) nucleus of the thalamus and in layer IV of primary somatosensory cortex (S1) in anesthetized rodents. Recordings were targeted to topographically aligned barreloids in VPm and barrel column in S1. Weak stimulation was applied to the whisker corresponding to the recorded barreloid/barrel column to elicit non-synchronous spiking. Putative monosynaptic connection between pairs of neurons were inferred using cross-correlation analysis. **B.** Experimental procedures used to establish paired recordings involve (1) Animal preparation includes surgeries. (2) S1 mapping, (3) identification of the whisker corresponding to the recorded barreloid, (4) targeting corresponding S1 barrel column and layer IV, (5) data collection for assessing monosynaptic connectivity by generating spikes via whisker stimulation, repeating step (3), (4) and (5) for recording additional pairs and (6) histology.

### Experimental workflow to establish paired recordings

Paired recordings in topographically aligned feedforward sensory regions in-vivo have been previously shown to be experimentally tractable (Bruno and Sakmann 2006; Bruno and Simons 2002; Reid and Alonso 1995; Swadlow and Gusev 2001; Wang et al. 2010) yet it remains a challenging process, and to our knowledge, detailed reports on experimental approaches have not been published in full. Here, we documented the steps in details. Figure 1B summarizes the general workflow for establishing and analyzing paired recording: 1) mapping primary somatosensory cortex (S1), 2) localizing the ventral posteromedial (VPm) nucleus of the thalamus, 3) achieving and verifying topographical alignment of recording electrodes across VPm and S1, 4) connectivity assessment through statistical analysis of the measured spiking activity from pairs of single-units across recording sites, and 5) histological verification of recording site locations.

Although the final electrode placement for paired recording involved thalamic electrode placement followed by placement of the cortical electrode, we found that an initial somatotopic mapping of cortex was critical for efficiently achieving topographical alignment. As part of the approach, we thus first performed coarse cortical mapping prior to thalamic localization. In rat, we employed primarily electrophysiological mapping approaches for this coarse mapping of S1. We targeted S1 using approximate stereotaxic coordinates and inserted a single electrode to obtain the functional location of several “barrels” using bregma as a reference point. Once we located the S1 region, we identified the stereotaxic location of three barrel columns containing neurons that were responsive to the movement of single primary whiskers (See Supplementary Figure 1 - https://doi.org/10.6084/m9.figshare.14393528.v1). We used this relative distance between columns/barrels to estimate the overall topography of S1 by overlaying a barrel map template scaled to fit the three data points. In mouse, taking advantage of the optical properties of the mouse skull (i.e. that it is relatively translucent), we employed intrinsic optical signal imaging (IOS) for the coarse cortical mapping. Intrinsic optical signals were acquired in response to separate, punctate deflections of three different single vibrissae. The corresponding cortical regions of activation were co-registered with the anatomy of the blood vessels and further used as a landmark for electrode placement. This triangulation methodology was adapted from previously published method from our laboratory (Gollnick et al. 2016; Millard and Stanley 2013). An example of this is shown in Figure 2A (see also Supplementary Figure 1A). The three images on the top row were acquired in response to punctate deflections of the Beta whisker (wBeta), the C2 whisker (wC2), and the B2 whisker (wB2), respectively. Each image represents the mean 0- to 6-s post-stimulus response (baseline subtracted and scaled) to a 4-s 10-Hz 1000 deg/s pulsatile stimulus, with the region of cortical activation appearing as a dark spot near the center of the image, at different locations for each whisker. We then fitted a barrel map template, recovered from histological brain sections from previous experiments onto the overall optical image of the brain surface through the thinned skull (template shown in bottom left of Figure 2A). The three centroids of the intrinsic imaging signals were used as reference points (Figure 2A, bottom middle) for transformation of the barrel map template involving scaling, rotation, translation and shearing. This transformed map along with the blood vessel image (Figure 2A, bottom right) was then used as a navigation guide for electrode placement for targeting a desired barrel column.

**Figure 2:**
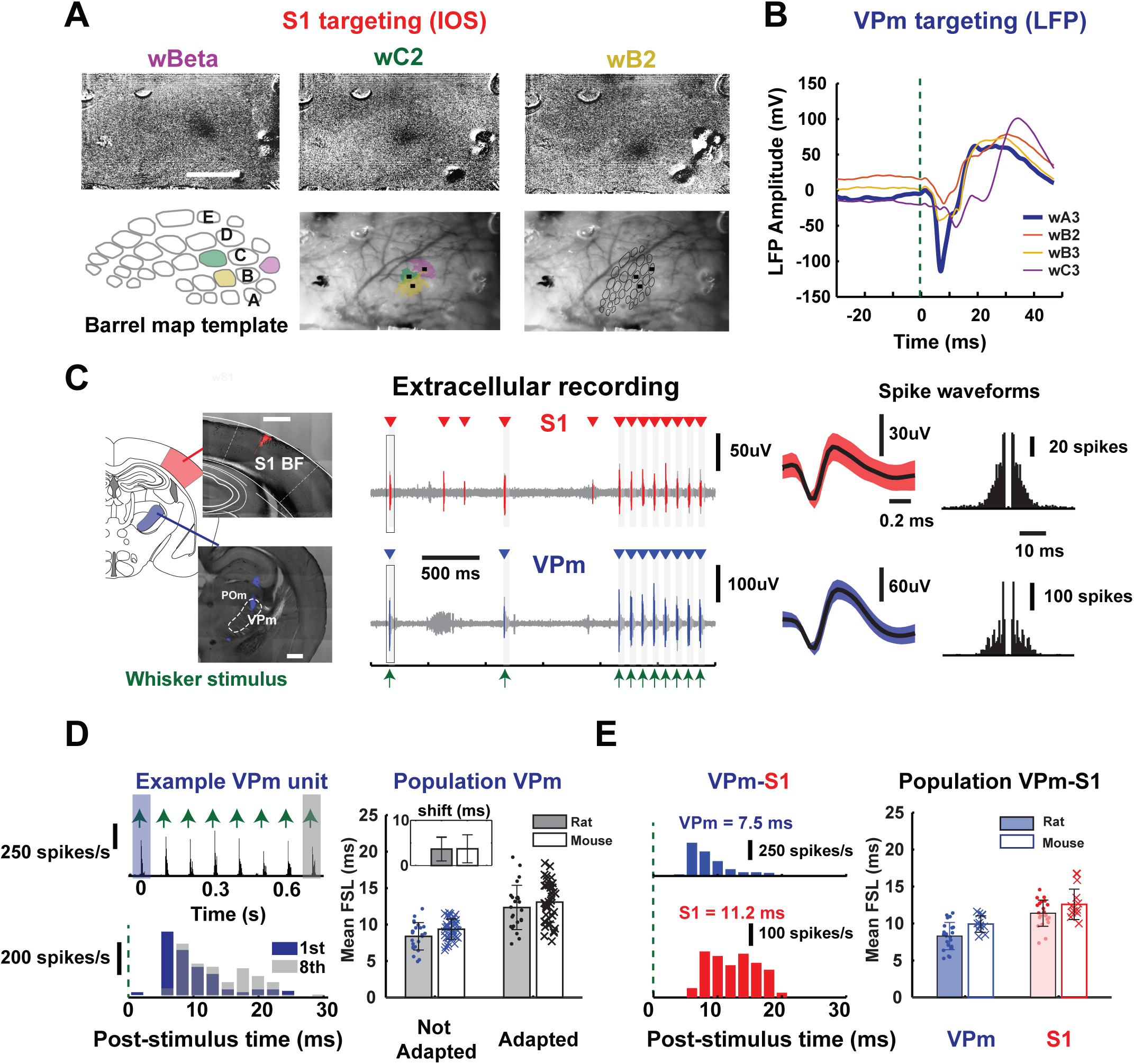
Targeting topographical alignment in vivo. **A.** S1 targeting: Location of S1 barrels was determined using intrinsic imaging (IOS) in mice. The three images on the top are images acquired in response to separate single punctate deflections of the Beta whisker, the C2 whisker, and the B2 whisker, respectively. A barrel template was compared to the optical image of the brain surface through the thinned skull and fitted based on the location of the centroid of barrels to provide guidance for electrode placement. **B.** (Left) VPm targeting: example mean LFP amplitude of evoked response (shown by downward deflection after stimulus onset) for four stimulated whiskers in mice. Note the largest and fastest response to the A3 vibrissa, and the comparatively weak responses to the adjacent B2, B3, and C3 vibrissae, lending support for VPm localization. **C. Left:** Histological slice showed electrode tracks marked with fluorescent dyes (see Method: Post-mortem histology) on coronal brain sections targeting VPm (bottom) and S1 layer IV (top) in a mouse. **Middle:** Example raw voltage traces from extracellular recordings performed simultaneously in VPm and S1 layer IV during sensory stimulation in a rat. Mean waveforms of isolated single units from each recording site were shown (shaded region indicates one standard deviation of spike amplitude), along with spike auto-correlograms of each unit in a rat (**Right**). **D. Left:** Example adapting response of a single thalamic unit from a rat, showing mean first-spike latency in response to first and last pulse of a 8Hz pulsatile, ongoing stimulus (1^st^ pulse: 5.18 ms, last pulse: 10.7 ms). **Right:** Population adapting response for thalamic units recorded in rats and mice (1^st^ pulse (rats): 8.39 ± 1.87 ms, last pulse (rats): 12.3 ± 3.04 ms, n = 24 neurons, N = 12 rats, 1^st^ pulse (mice): 9.37 ± 1.39 ms, last pulse (mice): 13.1 ± 2.78 ms, n = 39 neurons, N = 4 mice). **Inset:** All thalamic units that we recorded showed latency shift less than 20 ms (latency shift (rats) = 3.65 ± 2.62 ms, n = 24 neurons, N = 12 rats, latency shift (mice) = 3.71 ± 3.11 ms, n = 39 neurons, N = 4 mice), suggested that they were thalamic VPm units. **E. Left:** Example peri-stimulus time histogram (PSTH, 2 ms binsize) for a thalamic and cortical unit in the 30-ms window following punctate (600°/s) whisker stimulus (indicated by green dotted line at t=0). We computed mean first spike latency (FSL), defined as average latency of first spike in 30 ms response windows after stimulus presentation for each unit (VPm: 7.5 ms, S1: 11.2 ms). **Right:** Population mean first spike latency for all simultaneously recorded thalamic and cortical units in rats and mice (VPm (rats) : 8.30 ms ±1.84, S1 (rats): 11.40 ±1.77 ms, n = 22 neurons, N = 12 rats; VPm (mice): 9.80 ± 1.31 ms, n = 9 neuron, S1 (mice): 12.6 ± 2.06 ms, n =11 neuron, N = 1 mouse).

Once the cortex was coarsely mapped using the above approaches, we inserted an electrode to localize VPm (see Method: Electrophysiology). Specifically, we identified specific thalamic barreloids by manually deflecting each of the facial vibrissae individually in order to find an isolated unit that was maximally responsive to a single whisker only. Note that the procedure was nearly identical for rat and mouse, except that for the mouse, a silicon multi-electrode probe was utilized, and localization was performed using one of the probe sites, usually the deepest probe contact. Given that the probe was spanning several hundred microns on the thalamic recording site, we typically recorded from one to three barreloids simultaneously with two or more recording sites within a single barreloid. Final placement of the electrode was fixed when well-isolated units were detected on probe contacts. For verification of barreloid targeting within VPm, we compared local field potential (LFP) responses to individual stimulation of multiple nearby vibrissae. Figure 2B shows an example of LFPs recorded in the A3 barreloid of thalamus on one of the probe contacts. Note that the evoked LFP response exhibits an initial negative peak for the whisker A3 deflection (wA3, blue) that is substantially larger than responses to the deflection of other whiskers.

A practical challenge lies in the positioning of the recording electrodes to reach the multiple target areas. In a small subset of experiments, we optimized the electrode insertion angle by coating the electrodes with fluorescent dye (DiI) to mark the electrode track for post-experiment histological analysis. A range of insertion angles was tested for both S1 and VPm in mouse and rat. We found that the optimal electrode angle to target VPm thalamus in both rats and mice was 0 degrees from vertical (perpendicular to the brain surface) (Figure 2C left and Supplementary Figure 1B) For S1 targeting, we found that the optimal angle for rats was approximately 40-45 degrees from vertical and was approximately 30-35 degrees from vertical for mice for electrode penetration parallel to the cortical column/barrel, important for accurate targeting of specific cortical depths. The target recording site depths were determined based on published anatomical locations of the target structures (see Methods). Figure 2C shows typical electrode tracks targeting S1 and VPm in a mouse (left), along with the corresponding raw extracellular recordings from one VPm and one S1 electrode recording site (middle). Both the VPm and S1 extracellular recordings show sensory driven responses to deflections of the same whisker, the pattern of which is shown in the extracellular recordings of Figure 2C. The times of whisker stimulation are denoted with the green arrows, representing a pattern of two isolated whisker deflections, followed by an 8 Hz train of whisker deflections. The raw electrophysiological recordings were subsequently sorted into single-unit data based on conventional spike sorting approaches and only well-isolated units were retained for analysis (see Methods). For each recording, the identified putative single unit spiking (red, blue) is superimposed on the raw recording (light gray), and the corresponding spike times are denoted above each trace (red and blue triangles). For each case, the spike waveform and spike auto-correlogram are shown to the right in Figure 2C. A summary of the waveform size and isolation quality for all recorded VPm and S1 units is shown in Supplementary Figure 1C.

### Sensory response in topographically aligned VPm – S1 layer 4 regions

Achieving topographically aligned recordings across corresponding thalamic barreloids and cortical barrel columns necessitates the accurate targeting of the thalamic recording electrode/probe to the VPm nucleus of the thalamus. In particular, it is key to be able to distinguish whisker-responsive units located in VPm from those located in the adjacent posteromedial nucleus (POm) of the thalamus. In previous studies, POm units have been shown to exhibit lower evoked firing rate (Ahissar et al. 2000; Diamond et al. 1992; Landisman and Connors 2007; Sosnik et al. 2001) and much broader receptive fields than VPm units, responding to approximately six vibrissae on average, ranging from three to twelve whiskers (Castejon et al. 2016; Diamond et al. 1992). Hence, to ensure recordings from primarily VPm neurons, we selected thalamic units that showed restricted whisker sensitivity, usually only to one principal whisker (as previously shown in Figure 2B, VPm targeting (LFP)). The thalamic single units kept for subsequent connectivity analysis exhibited strong, reliable response (response reliability: 52.6 ± 24.62 %, not shown) to a punctate whisker stimulus with short latency (1^st^ pulse (rats): 8.39 ± 1.87 ms (mean ± SEM), last pulse (rats): 12.3 ± 3.04 ms (mean ± SEM), n = 24 neurons, N = 12 rats, 1^st^ pulse (mice): 9.37 ± 1.39 ms (mean ± SEM), last pulse (mice): 13.1 ± 2.78 ms (mean ± SEM), n = 39 neurons, N = 4 mice, Figures 2D and 2E, Supplementary Figure 2A). As an additional criterion, we measured the adapting properties of the thalamic units in response to repetitive, periodic whisker stimuli (Ahissar et al. 2000; Masri et al. 2008) (Sitnikova and Raevskii 2010). Previous studies have shown that neurons in POm exhibit dramatic adaptation to persistent sensory stimulation, where most Pom neurons failed to exhibit a response to stimuli ≥ 11Hz (Masri et al. 2008), and a significant shift in response latency with the adaptation, as compared to VPm neurons (Ahissar et al. 2000; Sosnik et al. 2001). Here, we showed a representative thalamic unit spiking response to an 8 Hz adapting whisker stimulus on the left in Figure 2D, with a PSTH in response to the full stimulus train (top) and a superposition of the stimulus evoked response to the 1^st^ (blue) and 8^th^ (gray) deflection in the train (bottom). This particular example shows strong stimulus-locked responses and moderate adaptation (i.e. reduction in response amplitude), with a relatively small increase in latency of spiking. In fact, all the thalamic units that we considered for further analysis showed only moderate spike frequency adaptation, with a maximal increase in latency of 8ms when comparing responses to the first and last stimulus of the train (Figure 2D Right, mean first spike latency (FSL), latency shift from Not-Adapted to Adapted = 4.28 ms ± 2.32 ms (mean ± SEM), n=21 cells from rat; 3.71 ± 3.11 ms (mean ± SEM), n = 39 neurons, N = 4 mice, Figure 2D right, inset). Furthermore, the response latency of thalamic neurons measured here is consistent with the latency of VPm neurons (peak latency <10ms) identified through a genetic validation approach (Wright et al. 2021). Note that there remains a small possibility that a subset of the thalamic units is POm in origin, and project to S1 inter-barrel areas (septa). However, this is unlikely to be the case for the thalamocortical pairs that showed monosynaptic connectivity, given that these thalamic and cortical units demonstrated mostly single whisker receptive fields (Furuta et al. 2009) and the likelihood of detecting synaptic connections in POm-S1 inter-barrel areas is much lower. Finally, a recent study showed that POm cells remain largely inactive (close to zero spontaneous firing rate) under isoflurane anesthesia (Zhang and Bruno 2019), making them extremely difficult to locate and record from under the conditions of this study, which when combined with the other factors described above make it unlikely that any of the recordings here are POm in origin. Again, note that the challenges here are somewhat specific to this particular brain region, but achieving definitive recordings in other brain regions would likely have similar challenges to those described here.

Previous studies have shown that the convergence of thalamic inputs onto topographically aligned cortical layer 4 neurons is generally relatively high, but with the probability of contacting regular spiking units (RSUs) much lower than fast spiking units (FSUs) (Bruno and Simons 2002; Swadlow 2003). We recorded from well-isolated cortical layer 4 neurons (see Supplementary Figure 1C) at a cortical depth taken from micromanipulator readings (Rat: 803.28 ± 175.28 um, mean ± SEM, n=21, data not shown; Mouse: 350-700 um, n =11, see Figure 5B). All cortical units included in this study were putatively layer 4 neurons. The mean first spike latency of cortical units in rat was 11.4 ms ± 1.77 ms (mean ± SEM, n= 22) and 12.6 ± 2.06 ms (mean ± SEM, n= 11) for mouse. We found that the relative latency difference between VPm and S1 layer 4 in aligned regions was comparable to the expected synaptic delay between VPm and S1 layer 4 (differences between VPm-S1 were ∼3 ms for both rat and mouse). An example of the response to a punctate whisker stimulus for a pair of VPm and S1 FS units with mean first-spike latency of 7.5 ms and 11.2 ms is shown on the left of Figure 2E.

### Inferring connectivity in the context of a signal detection framework

In general, neurons in the central nervous system require the concerted action of a relatively large number of pre-synaptic inputs to produce an action potential (Bruno and Sakmann 2006). Thus, the relationship between pre- and post-synaptic neurons is tenuous at best, reflected in often a very subtle increase in the probability of firing of the post-synaptic neuron a few milliseconds after the firing of a pre-synaptic neuron. The analysis of spike trains from a pair of neurons can thus be utilized for a simple binary classification of a neuronal pair either being ‘not connected’ or ‘connected’ (Figure 3A).

**Figure 3:**
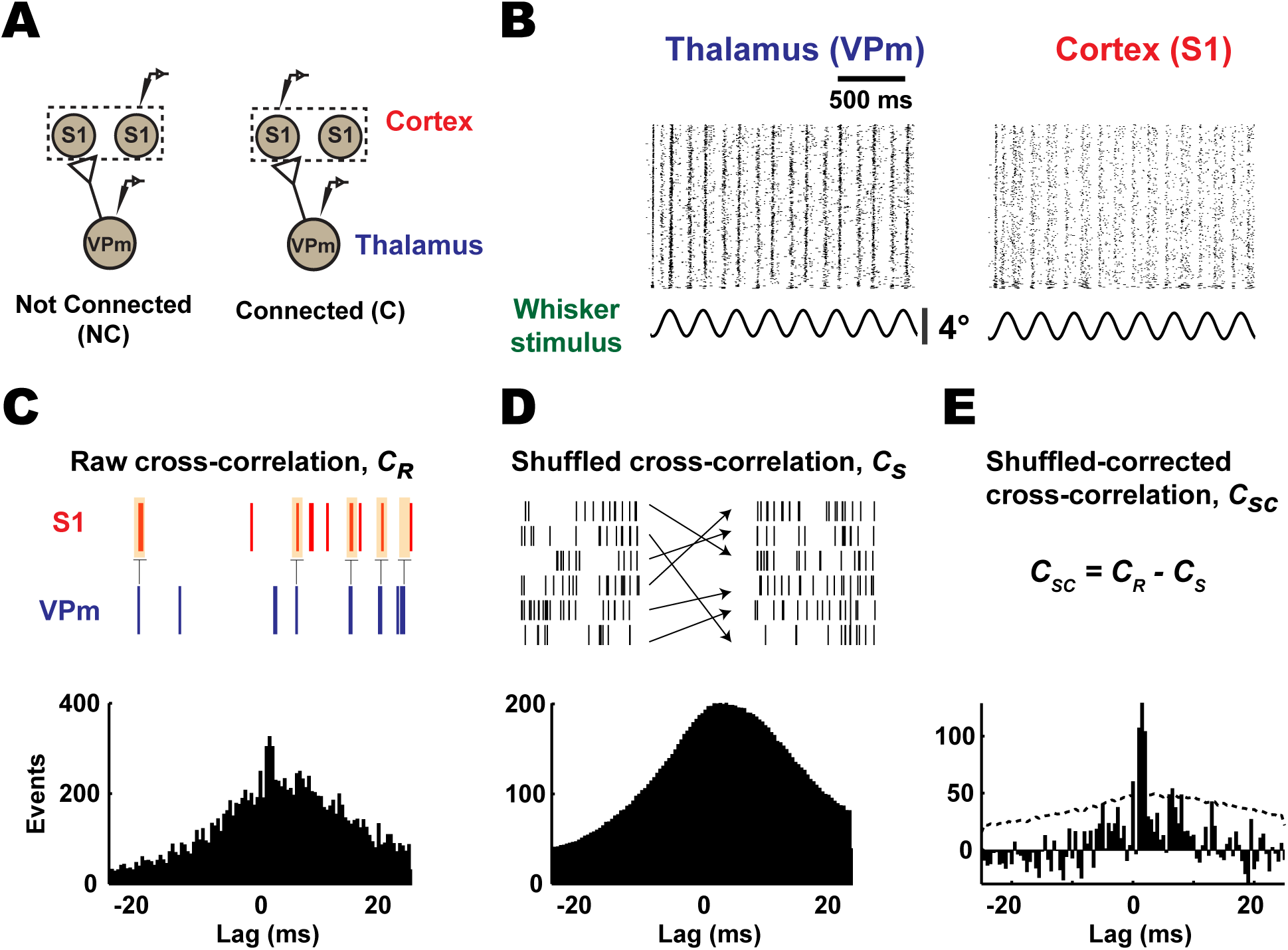
Monosynaptic connectivity inference using cross-correlation analysis. **A.** Inferring monosynaptic connectivity from extracellular recordings performed in topographically aligned thalamocortical regions in-vivo resulted in binary consequences. A pair of neurons can be putatively classified as ‘connected’ or ‘not connected’, as shown in the schematic. **B**. Raster plots showing whisker evoked spiking response under low velocity sinusoidal stimulation (mean velocity: 25 deg/s) for a representative example from thalamus and cortex in a rat. **C**. All cross correlograms were computed using VPm spike train as a reference. Occurrences of cortical spikes were measured at various time lags (25 ms window before and after a thalamic spike, with 0.5 ms step size; see Method: Monosynaptic Connectivity). **D.** Stimulus-driven cross-correlograms were constructed between the original reference VPm spike trains and the trial-shuffled cortical spike trains. **E.** Shuffled-corrected cross-correlograms were generated by subtracting the mean of shuffled cross correlograms (averaged from 1000 iterations) from the raw cross-correlograms. Dotted gray lines denoted 3.5 standard deviation of the shuffled distribution.

In order to infer connectivity, for each recorded pair of neurons, the raw spike cross-correlogram is calculated using standard approaches (C_r_, Figure 3C) (see Methods), effectively yielding a histogram of cortical firing relative to thalamic spike times. Because the analysis is based on spiking activity, and the baseline spiking activity can often be relatively low, we drove the thalamocortical circuit in-vivo with a weak whisker stimulus (Figure 3B, sinusoidal deflection, 4 Hz, mean velocity: 25 °/s), which is known to enhance firing rates with minimal impact on firing synchrony across neurons (Bruno and Sakmann 2006). To correct for correlated stimulus-locked activity, we generated a shuffled-corrected spike cross-correlogram (C_sc_, Figure 3E) by subtracting the trial shuffled spike cross-correlogram (C_s_, Figure 3D) from the raw spike cross-correlogram. The “shuffled cross-correlogram” is generated using the same procedure used for the raw cross-correlogram, except that the trials of the thalamic and cortical spiking activity are randomized relative to each other. This effectively destroys any elements of the cross-correlogram that are not due to the stimulus. Shown in this example is the qualitative signature of monosynaptic connectivity – a prominent peak in the shuffled-corrected cross-correlogram for small positive lags that would be consistent with a single synaptic delay.

Then, to conclude that a neuronal pair was ‘connected’, we adopted two criteria that expanded from previous studies (Bruno and Sakmann 2006; Reid and Alonso 1995; Swadlow and Gusev 2001) based on both the raw and shuffled-corrected cross-correlogram: (Criterion 1) a notable sharp, millisecond-fine peak is observed within a narrow lag of 1-4 ms after a thalamic spike, and (Criterion 2) this fast ‘monosynaptic peak’ is significant or still present after accounting for (subtracting) stimulus-induced correlation. Note that a peak is defined as the bin in the raw cross-correlogram (0.5 ms bin size) that contains the maximum number of events. In order to fulfill both criteria, we required that the peak detected in the 1-4 ms range to have the largest correlation out of all the bins in the range of +/- 25 ms window and this correlation is significant (>3.5 SD) against shuffled data. For the example ‘connected’ pair in Figures 3C-E and a separate ‘not connected’ pair, Criterion 1 is evaluated from the raw cross-correlation as shown in Figure 4A. For the ‘not connected’ example (raw cross-correlogram on the left), a peak was detected outside the central 1-4ms lag, and thus fails Criterion 1. The example ‘connected’ pair, the raw cross-correlogram on the right, exhibits a peak within the central 1-4 ms lag (shown as a vertical gray band), and thus passes Criterion 1. Figure 4A illustrates how Criterion 2 is estimated from the shuffled-corrected cross-correlation. We evaluate the prominence of the peak relative to a distribution of peak magnitudes (same bin with maximum number of events) from the shuffled cross-correlograms (1000 iterations). Specifically, we define the metric related to Criterion 2 as the peak height, h, computed using the maximum of the C_sc_ (0.5 ms bin size) within 1-4 ms lags, normalized by the standard deviation of the shuffled cross-correlogram. This metric is thus the number of standard deviations the peak of the shuffled-corrected cross-correlogram within the 1-4ms lag is above the shuffled distribution. This is shown in more detail for this example in the inset in the bottom row of Figure 4A, where the central portions of the shuffled-corrected cross-correlograms are shown for each case. In order to pass Criterion 2, the peak height of the shuffled-corrected cross-correlograms must be greater than 3.5 standard deviations of the shuffled data, which corresponds to a 99.9% confidence interval for each bin.

**Figure 4:**
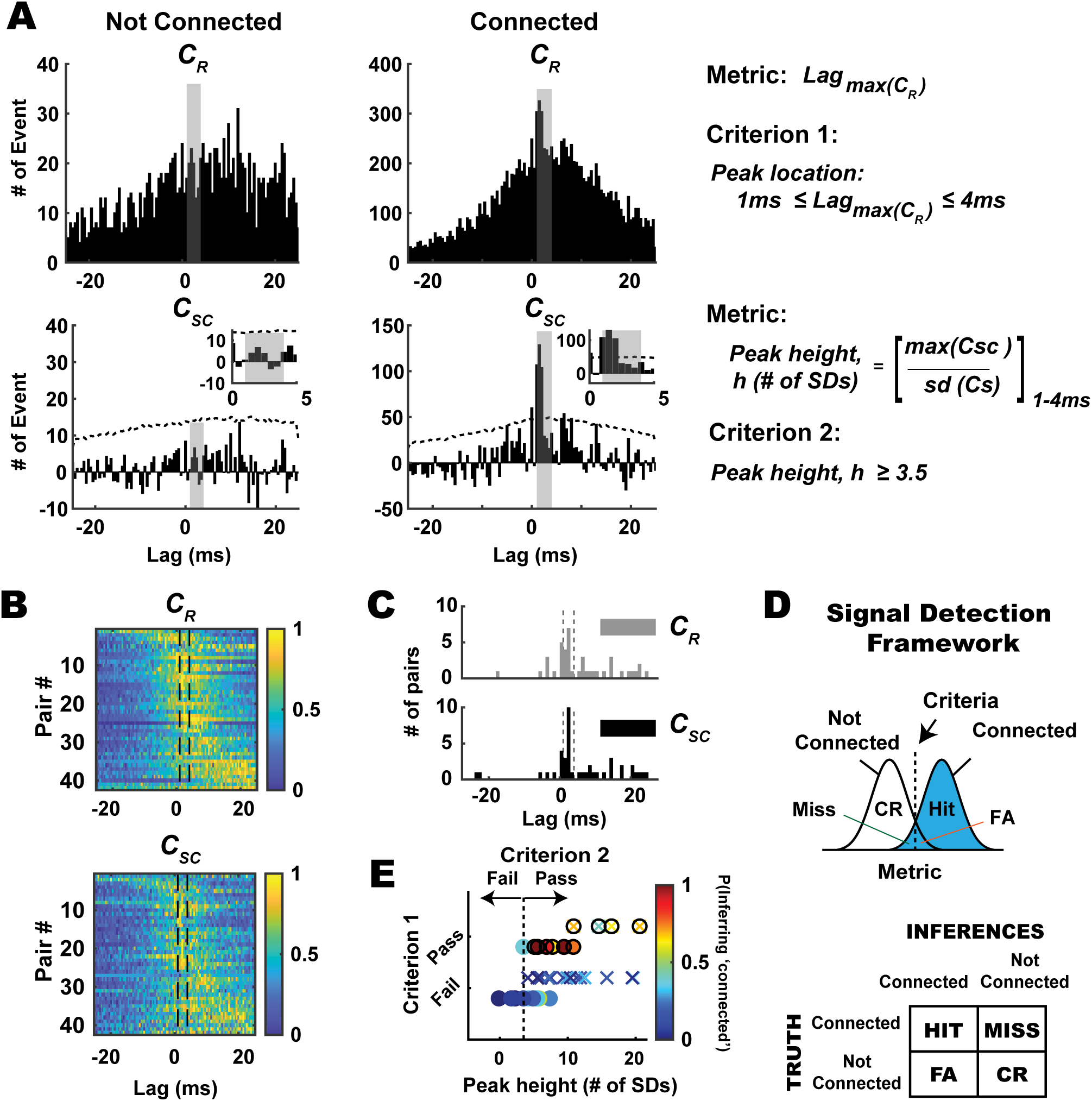
Evaluation of monosynaptic connection inference in the context of a signal detection framework. **A.** In the context of signal detection framework, we defined two distinct metrics and criteria to classify neuronal pairs into ‘connected’ and ‘not-connected’ distribution. Showing here are representative example pairs from each condition. (Top) Note the qualitative difference in raw cross correlograms (C_R_), one with broad and distributed spikes in cross-correlogram (left) and another with sharp peaks (right) in 1-4 ms time lags (shaded gray). The first metric is the maximum peak of the raw cross-correlogram (C_R_), and Criterion 1 was fulfilled if the maximum peak of the raw cross-correlograms was within 1-4 ms lag. (Bottom) After corrected for stimulus-driven correlations, we further quantified the significance of the peaks detected in raw cross-correlogram. Hence, second metric used here is the peak height within the window of interest (1-4 ms bin), measured as maximum peak value in shuffled-corrected cross-correlogram (C_SC_), normalized to the number of standard deviations (# of SDs) with respect to the shuffled distribution. Criterion 2 was fulfilled if the peak exceeded 3.5 standard deviation of the shuffled distribution. **B.** As expected, the majority of simultaneously recorded thalamocortical pairs exhibited peak correlation after 0 time lag (n = 42 pairs, rats – 22; mice −20). Histograms show the number of events in each bin of raw cross-correlograms (top) and shuffled-corrected cross-correlograms (bottom), sorted by latency of maximum peak in the cross-correlograms. Data were normalized to maximum peak and the colors in each row show the number of events for an individual pair, normalized to maximum peak. **C.** Distribution of the lags of the peak location in the raw and shuffled-corrected correlograms across all recorded pairs. Note that the peak locations for raw and shuffled-corrected correlograms were largely the same, shifted only by 1-2 bin size (0.5ms bin). **D.** Within this framework, four possible outcomes are possible. Hit: Connected and inferred to be connected. Miss: Connected but inferred to be not-connected. False Alarm: Not connected but inferred to be connected. Correct Rejection: Not-connected and inferred to be not-connected. **E**. In this context, we found that 11/42 pairs have putative monosynaptic connection (denoted as solid circles), 31/22 pairs were not connected. (n = 22 neurons, N = 12 rats; n = 20 neurons, N = 1 mouse). Note that Criterion 1 was a binary classification – pairs having peak locations from their raw cross-correlograms within the 1-4ms lag pass Criterion 1, and those which do not, fail Criterion 1. For Criterion 2, the dashed vertical line represents the 3.5 SD for the peak height metric – pairs having peak height above this criterion line pass Criterion 2. Only pairs that passed both Criterion 1 and 2 in Figure 4D (bottom) were classified as ‘connected’ (anything to the right of the vertical dashed line and on the top row) and the rest were classified as ‘not connected’. For each recorded pair, probability of inferring monosynaptic connection (i.e. probability of bootstrapped data satisfying Criterion 1 and 2) was depicted with a color bar. Note that 1000 iterations were performed for each data set.

Thus, we conclude that the shuffled-corrected cross-correlograms on the left fails Criterion 2 as the peak falls below the criterion line (3.5 SD, depicted as dashed line) whereas the shuffled-corrected cross-correlograms on the right passes Criterion 2. Note that for the “not connected” pair shown in Figure 4A, the shuffled-corrected cross-correlogram also revealed the presence of the global peak outside the 1-4 ms central lag, similar to that of the raw cross-correlogram. Thus, it is possible to utilize the shuffled-corrected cross-correlogram for the evaluation of Criterion 1 in some cases, but overall, we found that Criterion 1 was more robustly evaluated utilizing the raw cross-correlogram. The raw and shuffled-corrected correlograms for all recorded thalamocortical pairs are shown in Figure 4B - normalized to the peak in each correlogram, ordered from earliest to latest peak, top to bottom. The vertical lines in these plots highlight the 1-4ms lag range described in Figure 4. Figure 4C shows the distribution of the lags of the peak location in the raw and shuffled-corrected correlograms across all recorded pairs. Overall, the peak locations for raw and shuffled-corrected correlograms were similar, shifted only by 1-2 bin size (0.5ms). For all the monosynaptically connected pairs in this paper, the mean peak location was 1.18 ± 1.08 ms (mean ± SEM, n = 11), and with a median of 1.5 ± 0.89 ms.

Note that to be classified as connected, a candidate pair of neurons must pass both Criterion 1 and Criterion 2. Using these metrics and criteria, we can consider the result of cross-correlation analysis for each pair as an inference problem in the context of a signal detection framework, where we conceptualize the distribution of metric values for the non-connected pairs as “noise”, and for the connected pairs, as “signal”. Applying the criteria on the metric values yields four possible outcomes: Hit (connected pair classified/inferred as such), Miss (connected pair classified/inferred as ‘not connected’), False Alarm (not-connected pair classified/inferred as ‘connected’), and Correct Reject (not-connected pair classified/inferred as such) (Figure 4D). With these metrics and criteria, we classified measured neuronal pairs into ‘connected’ and ‘not-connected’ distributions, with 11/42 pairs having a putative monosynaptic connection and 31/42 pairs having no apparent connection (Figure 4D bottom, Rat: n = 22 pairs, Mouse = 20 pairs). For Criterion 1, this was a binary classification due to biological constraints (single synaptic delay) – pairs having peak locations from their raw cross-correlograms within the 1-4ms lag pass Criterion 1, and those which do not, fail Criterion 1. For Criterion 2, the dashed vertical line represents the 3.5 SD for the peak height metric – pairs having peak height above this criterion line pass Criterion 2. Only pairs that passed both Criterion 1 and 2 in Figure 4D (bottom) were classified as ‘connected’ (anything to the right of vertical dashed line and on the top row) and the rest were classified as ‘not connected’. Note that while this is a binary classification, and the end-result is a labeling of a pair as ‘connected’ or ‘not-connected’, not all classifications are equivalent, with some having substantially more confidence in the classification than others. By utilizing a bootstrapping approach (see Methods) and evaluating the likelihood of specific outcomes using the pre-defined metrics and criteria for each recorded pair, we were able to attach a probability measure (Figure 4E bottom, color bar, P(Inferring ‘connected’)) along with each connectivity inference. In Figure 4E bottom, some of the cases that passed Criterion 1 have peak heights that are substantially far away from the line for Criterion 2. As expected, the likelihood of inferring monosynaptic connection for these cases was higher (∼0.6-1) and thus we have more confidence in the assertion, as compared to cases that are very close to the criterion line for Criterion 2, or failed Criterion 1.

### Probabilistic measure of connectivity inference across brain structures in large-scale recordings

Through the advent of high channel-count electrophysiological recording techniques (Chung et al. 2019; Jun et al. 2017; Rios et al. 2016), the diversity of recording quality, cell type, and possibilities for connectivity has expanded tremendously. To demonstrate how our framework can be beneficial, Figure 5 shows an example of simultaneous topographically aligned recordings from silicon multi-electrode probes inserted in VPm thalamus (Figure 5A) and S1 (Figure 5B). Specifically, focusing on portions of the probes that were assessed to be topographically aligned, a subset of channels from the VPm probe (Channels A-D) yielded 5 thalamic neurons (VPm 1-5), while a subset of channels from the S1 probe (A-D) yielded 4 cortical neurons (S1 1-4). These neurons were recorded during weak sinusoidal (25 deg/s, 4Hz) whisker stimulation. In this multi-dimensional analysis, each of the 20 thalamocortical pairs were assessed for the possibility of connectivity. Table 1 in Figure 5C shows the matrix of binary outcomes of monosynaptic connectivity inference from cross-correlation analysis of the full dataset across the recording sites. Out of the 20 VPm-S1 pairs, 4 were judged to be connected according to our criteria. Note that while the connectivity from an individual barreloid to the homologous cortical barrel is relatively high (it has been estimated that approximately 1 in 3 VPm neurons within a barreloid is connected to a particular neuron in cortical layer 4 of the homologous barrel (Bruno and Sakmann 2006)), the relatively stringent criteria of the inference coupled with other factors such as the potential to record from multiple nearby VPm barreloids and variable single unit quality make this outcome typical. Shown in Figure 5C Table 2 are the probabilities of connectivity associated with each pair using the bootstrapping method (n = 1000 iterations) on the data (Table 2), as previously described. In general, the binary inference for a particular pair corresponded to the bootstrapped confidence levels – connected pairs had a relatively high probability of connectivity from the bootstrapping (see all ‘connected’ pairs in Supplementary Figure 4) while not connected pairs had a relatively low probability of connectivity from the bootstrapping (p<0.4). We found that the binary outcomes of monosynaptic connectivity inference could result in different ranges of probability of inferring monosynaptic connection (0.4-1). One factor that could affect this probability is data-length. Not surprisingly, we found that the ‘connected’ pair that yielded the lowest probability in this matrix (p = 0.41) had about 3900 spikes (in terms of geometric mean), as compared to the others (4900, 6900 and 10000 spikes).

**Figure 5:**
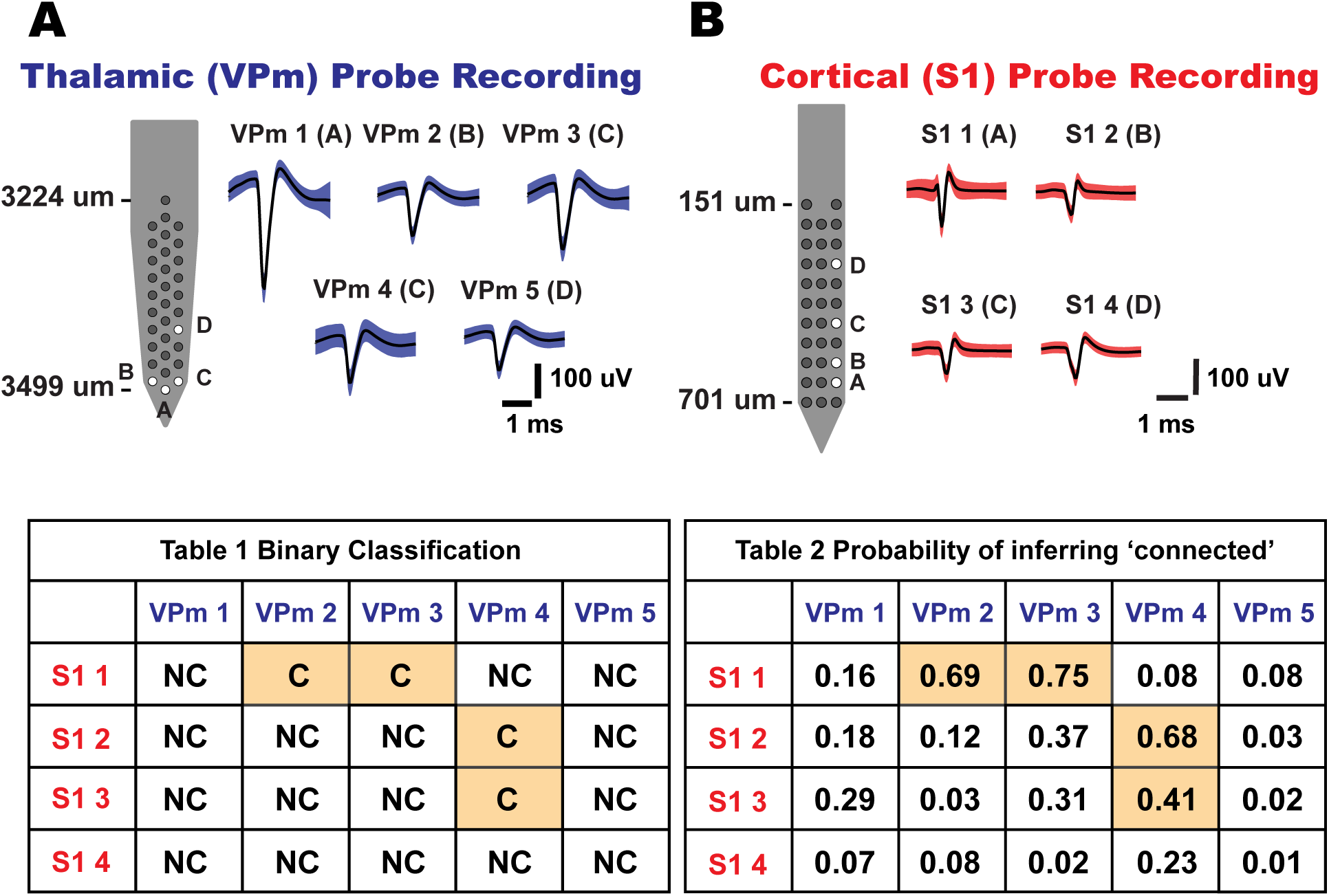
Connectivity matrix for topographically aligned, simultaneous multi-site recordings. **A.** Thalamic (VPm) probe recording. Five whisker-responsive thalamic units were isolated from a 32-channel silicon probe site labeled A-D. Mean waveforms of single units were shown on the right (shaded region indicates one standard deviation of spike amplitude) (n = 5 neurons). **B.** Cortical (S1) probe recording. Four whisker-responsive units, putatively from layer IV barrel cortex, were isolated from a 32-channel silicon probe site labeled A-D. Mean waveforms of units were shown on the right (shaded region indicates one standard deviation of spike amplitude) (n = 4 neurons). Table 1 shows binary outcomes of the monosynaptic connectivity inference based on Criterion 1 and 2 of cross-correlation analysis. Table 2 shows the connectivity matrix tabulating the probability of inferring a putative monosynaptic connection using the bootstrapping method for each thalamocortical pair in Table 1 (bootstrap iteration = 1000).

### Data-length effect on monosynaptic connection inference

As previously shown, inference of monosynaptic connectivity using cross-correlation analysis on extracellular signals is highly dependent on the amount of collected data. Based on existing literature, the recommended numbers of spikes were highly variable, ranging from 2000 to 10000 spikes (Swadlow and Gusev 2001; Wang et al. 2010), but were presented more as a “rule-of-thumb” than based on systematic evaluation. Here, we systematically evaluated data-length dependence effects on the connectivity inference outcomes. More specifically, we measured the data-length effect by performing bootstrapping on the full dataset by randomly selecting segments of data of increasing duration, as illustrated in Figure 6A. Note that for this analysis, the effects of data-length were evaluated in the context of Criterion 2, as this is the measure more prominently affected by data-length. In general, the analysis is primarily sensitive to the number of spikes used in the estimates, rather than the time duration of experimental data collection, and is sensitive to the number of both VPm and S1 spikes (i.e. sufficient spiking from both is requisite). For this reason, we utilized the geometric mean of the number of VPm and S1 spikes, calculated as the square root of the product of the number of VPm and S1 spikes. Because we did not have access to an established “ground truth” of a pair being either connected or not connected, we utilized example pairs in which the analysis revealed a very clear classification, which we subsequently utilized as ground truth for the analysis.

**Figure 6:**
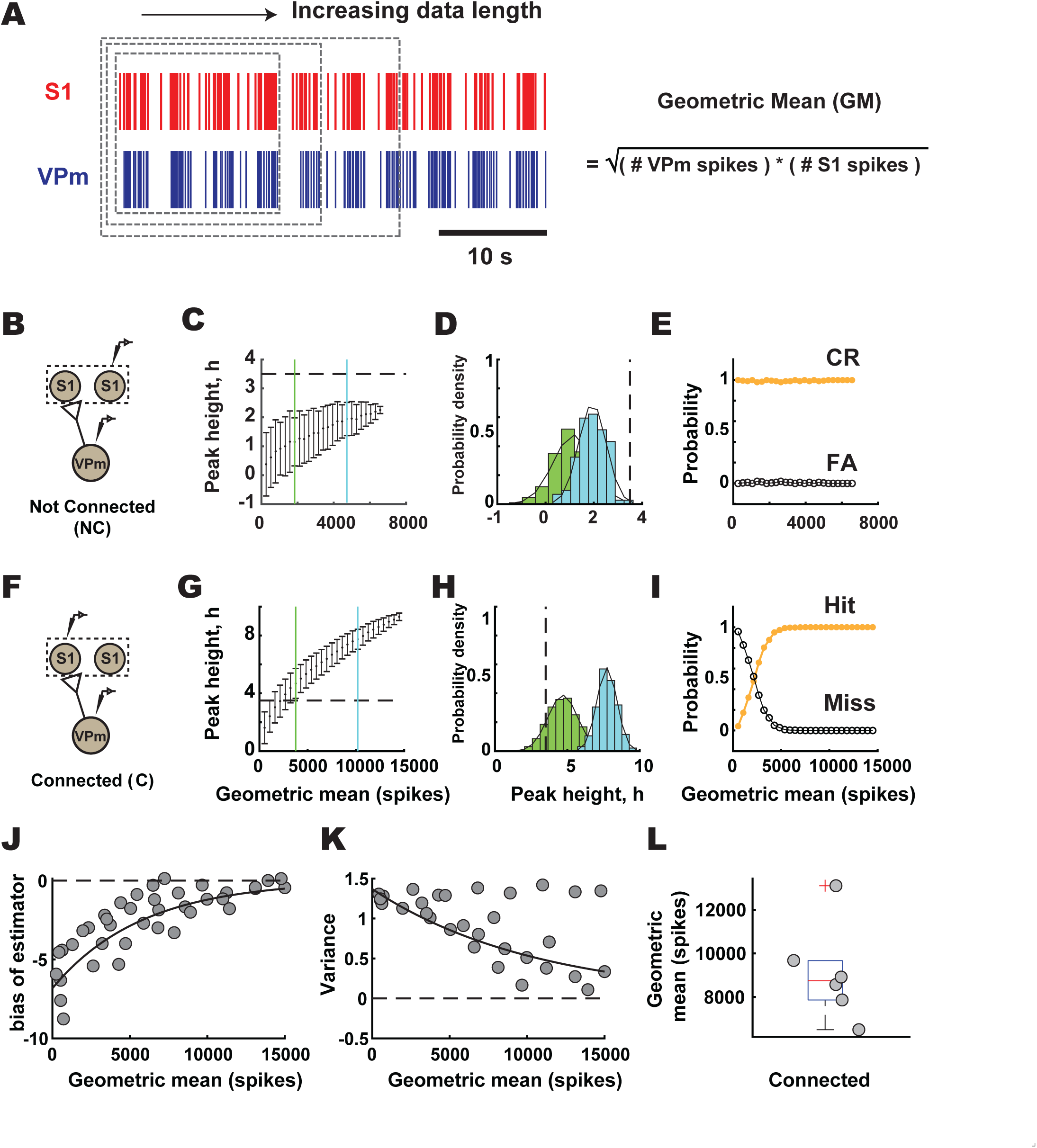
Data-length dependence effect on monosynaptic connection inference. **A.** Data-length dependence effect on connectivity inference was evaluated using a subsampling method. Data length was measured in terms of geometric spikes, which was calculated by taking the multiplication of the square roots of total VPm and S1 spikes. Random subsampling of the dataset was performed in the unit of trials with 1000 iterations for each condition. **B.** Representative example pair of neurons from ‘not connected’ distribution in a rat. **C**. Mean and standard deviation of peak height from cross-correlogram were computed for each subsample of ‘not-connected’ example. Blue line, short data-length condition (GM: 1837 spikes). Green, long data-length condition. **D**. Distribution of peak height after bootstrapping for two data-lengths labeled in part I, [Blue (GM): 1837 spikes; Green (GM): 4730 spikes]. **E**. Probability of outcome for this example, CR: Correct Reject; FA: false alarm. **F-I.** Similar to B-E but for ‘connected’ example in a rat [Blue (GM): 3760 spikes, hit rate: 87.7%, miss rate: 12.3%; Green (GM): 10201 spikes, hit and miss rate: 100%, 0%]. **J**. Bootstrap estimator of bias for each data length. Scatterplot of population data for connected pairs (n=6 pairs, N = 3 rats). Bold, exponential fit (R^2^=0.76). **K.** Variance of peak height at each data-length for pairs shown in J. Bold, 1^st^ order polynomial (R^2^ =0.62%). **L.** Geometric mean for all connected pairs (Median: 8741 spikes, n = 6 pairs, N = 3 rats).

Shown are the results from such an analysis for a ‘not connected’ pair (Figure 6B-E) and a ‘connected’ pair (Figure 6F-I). We found that, when a functional connection was obviously not present, the correlated firing activity between a pair of neurons was indistinguishable from that expected from stimulus-induced correlation, reflected in the metric remaining well below the criterion line (3.5 SD) for all values of data-length, while however steadily increasing as a function of the geometric mean (also accompanied by a decrease in the variability of the metric as reflected in the SEM) (Figure 6C). Figure 6D shows the distributions of the estimated connectivity metric h for two particular data-lengths (geometric means of 1837 spikes and 4730 spikes), relative to the criterion line. Both of these distributions are clearly to the left of the criterion line, corresponding to high correct reject and low false alarm rates, relatively unaffected by data-length (Figure 6E).

In contrast, when a functional connection was apparent, limited data confounded the inference of ‘connected’ as the metric remained below the criterion line for smaller data-lengths (Figure 6G). For two particular data-lengths (geometric means of 3760 and 10201 spikes), the distributions of the estimated connectivity metric h are shown in Figure 6H, relative to the criterion line. While the longer data-length resulted in a distribution of the peak height that was clearly above the criterion line (easily passing Criterion 2), the shorter data-length resulted in a distribution whose mean was above the criterion line (barely passing Criterion 2), but with a substantial portion of the distribution below the criterion line, resulting in a hit rate of 87.7% and miss rate of 12.3%. We found that the data-length required to reach a consistently correct inference was approximately 5000 spikes for this example (Figure 6I).

From the results in Figures 6B-E and 6F-I, it appears that data-length was critical only for correctly identifying a connected pair, and did not directly affect the inference related to a “not connected’ pair – in other words, the analysis is prone to type II errors as opposed to type I, and the type II error is strongly dependent upon data-length. It is important to point out that the data-length needed for sufficiently reducing the probability of error in the inference in Figure 6I is dependent upon the ground truth value for the connectivity metric h, as well as the variance in the estimator. We thus analyzed the estimator bias and variance across six ‘connected’ pairs, as a function of the data-length, which we could systematically vary by utilizing sub-sampling of the full datasets. Figure 6J shows the bias in the estimator for the connectivity metric h, as a function of the geometric mean number of spikes, with generally a negative bias (an underestimate) and a clear exponential decrease in bias with data-length as expected, as the full data-length is approached. Figure 6K shows the corresponding variance in the estimator of the connectivity metric h, again as a function of the geometric mean number of spikes. Although some of the measures exhibited an apparent invariance to the data-length, overall, this quantity decreased exponentially with increasing geometric mean number of spikes, again as expected. For the set of ‘connected’ pairs of thalamic and cortical neurons here, the geometric mean number of spikes required for the lower bound of resampled data to pass Criterion 2 is shown in Figure 6L, with a median of approximately 9000 spikes.

### Thalamic synchrony effects on monosynaptic connection inference

Without having access to the subthreshold activity of the postsynaptic cortical neurons, assessment of connectivity through the co-occurrences of spiking activity in thalamic and cortical neurons often necessitates activation of the intact circuitry with exogenous stimulation due to relatively low intrinsic spontaneous firing rates. The analysis of Figure 6 revealed a clear motivation for acquiring larger numbers of spikes, which can obviously be facilitated through increased mean firing rates. In sensory pathways, previous studies have used sensory stimuli of varying strengths to evoke higher firing rate in primary sensory areas (Bruno and Sakmann 2006; Reid and Alonso 1995; Sedigh-Sarvestani et al. 2017; Wang et al. 2010), citing a rule-of-thumb which involves increasing the number of spikes through external stimulation, but not with a stimulus so strong as to induce stimulus-driven synchronization in spiking. Although this approach is logical, it remains ad hoc, and the exact ramifications are not clear. Here, we utilized silicon multi-electrode probes in the thalamic VPm and S1 layer 4 to quantify changes in neural activity across spontaneous and stimulus driven conditions, and performed analyses to systematically evaluate the potential effects of stimulus-driven changes in firing rate and synchrony on the monosynaptic connection inference between VPm and S1 pairs. Figure 7A shows simultaneous recordings from two thalamic neurons (VPm1 and VPm2) within the same thalamic barreloid and two corresponding cortical S1 neurons in the homologous barrel column. For this example, VPm Unit 1 and S1 Unit 1 were revealed to be ‘connected’, while VPm Unit 2 and S1 Unit 2 were clearly ‘not connected’. Under increasingly stronger stimulus drive, from spontaneous (left), to weak-sinusoidal whisker stimulus drive (middle), and to strong repetitive, punctate whisker stimulus drive (right), there is a general increase in firing rate across the recorded cells in both VPm and S1 (Figure 7B). In addition to the modulation of mean firing rate, we found that increasing stimulus drive affected the inference of connectivity by producing inconsistent conclusions across stimulus conditions, especially obvious with increasingly strong stimuli. Interestingly, we found that this preferentially affected the location of the peak in the cross-correlogram, which is associated with Criterion 1 evaluated through the raw cross-correlogram. Figure 7C shows the raw and shuffled-corrected spike cross-correlograms across the stimulus conditions for the connected pair VPm Unit 1-S1 Unit 1, and Figure 7D shows this for the not-connected pair VPm Unit 2-S1 Unit 2. Note that we were able to maintain stable recordings for this particular group of neurons over a relatively long experimental time period, enabling us to collect a sufficient number of spikes in the spontaneous condition in order to make inferences regarding connectivity, which we can treat as ground truth here (VPm Unit 1-S1 Unit 1 connected, VPm Unit 2-S1 Unit 2 not-connected). First considering the connected pair VPm Unit 1-S1 Unit 1 (Figure 7C), we found that the inference remained consistent (‘connected’) for spontaneous (I) and sinusoidal (II) stimulus conditions as it passed both Criterion 1 and 2. However, for the repetitive transient (III) stimulus condition (bottom row), while the peak height in the shuffled-corrected cross-correlogram exceeded the criterion line (thus satisfying Criterion 2, 7C bottom panel), there was a disappearance of an isolated peak in the 1-4 ms lag of the raw cross-correlogram, which is a violation of Criterion 1. This led to the incorrect inference of ‘not connected’ for the case of the strong, transient stimulus, or a “miss” in the language of signal detection theory. In our analysis, we have 12 thalamocortical pairs for which we have all stimulus conditions, where we consider the spontaneous condition as the “ground truth”. Of these 12 pairs, 3 pairs were connected (C) and 9 not connected (NC), as determined from the spontaneous condition, which we consider “ground truth”. For the 9 NC pairs, 6 pairs were incorrectly classified as C in the strong (transient) stimulus case, and the remaining 3 pairs continued to be correctly classified as NC. For the 3 C pairs, 2 pairs were incorrectly classified as NC in the strong (transient) stimulus case, and the remaining 1 pair continued to be correctly classified as C. For this relatively small number of pairs, it is difficult to identify a particular pattern in this, but we can generally say that both types of errors can emerge from the strong stimulus that synchronizes the thalamic population, as we also demonstrate in the simulations.

**Figure 7:**
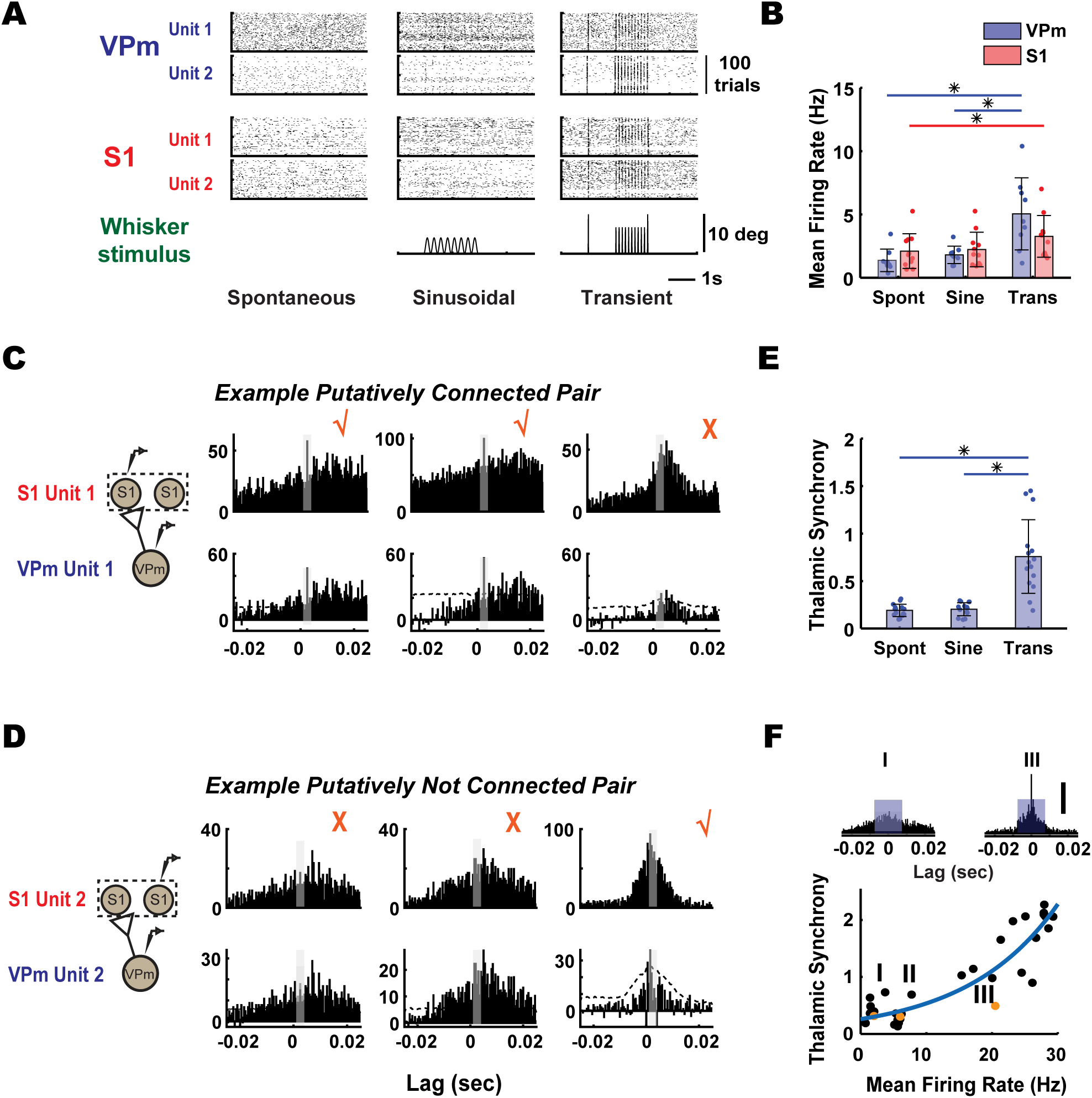
Thalamic synchrony effect on monosynaptic connection inference. **A.** Raster plots from simultaneously recorded thalamic and cortical units under spontaneous (no stimulus), sinusoidal stimulus (mean velocity: 25 deg/s) and transient inputs (1200 deg/s) in a mouse. **B.** Mean firing rate of thalamus (VPm, blue) and cortex (S1, red) of neuronal pairs were quantified across all three stimulus conditions. VPm: spontaneous: 1.38 ± 0.89 Hz, sinusoidal: 2.11 ± 1.37 Hz, transient: 5.05 ± 2.85 Hz, mean ± SEM (n = 9 neurons); S1: spontaneous: 2.11 ± 1.37 Hz, sinusoidal: 2.23 ± 1.36, transient: 3.27 ± 1.65 Hz (n = 11 neurons, N = 1 mouse) (* indicating p< 0.5, Wilcoxon signed-rank test with Bonferroni correction) **C.** Monosynaptic connectivity inference for one representative example of ‘connected’ thalamocortical pair from mouse. [Checkmark symbols indicated that the pair was being classified as putatively connected; cross symbols indicated that the pair was being classified as not connected]. **D.** Same as C, but for not-connected example. **E.** Thalamic synchrony was computed across three stimulus conditions, calculated as number of synchronous events between two thalamic units that occur within central window (± 7.5ms). (* indicating p<0.05, Wilcoxon signed-rank test, with Bonferroni correction). Spontaneous: 0.20 ± 0.07; Sinusoidal: 0.19 ± 0.06; Transient: 0.76 ± 0.39, mean ± SEM (n = 9 neurons, 15 pairs, N = 1 mouse). **F**. (Top) Raw spike cross-correlograms for the two VPm units shown in the top row of A in spontaneous and transient stimulus conditions. (I. Spontaneous, III. Transient). Blue box on each cross-correlogram represents the central window (± 7.5ms) used for thalamic synchrony computation. Scale bar represents 100 spikes in cross-correlograms. (Bottom) The relationship between thalamic synchrony and mean firing rate of respective thalamic pairs was quantified across various stimulus conditions, including spontaneous, sinusoidal, and transient stimuli with six different velocities (n = 5 neurons, 10 pairs, N = 1 mouse). Orange symbols represent thalamic synchrony across three stimulus conditions (I. Spontaneous, II. Sinusoidal, III. Transient). Blue line showing exponential fit of the relationship.

Turning to the ‘not-connected’ pair VPm Unit 2-S1 Unit 2 in Figure 7D, note that for the spontaneous (I) and sinusoidal (II) case, while the peak height of the shuffled-corrected cross-correlogram exceeded 3.5 SD (thus passing Criterion 2, not shown), we correctly inferred ‘not connected’ for these stimulus conditions given the violation of Criterion 1. Surprisingly, we incorrectly inferred ‘connected’ for the repetitive transient (III) stimulus condition as fast peaks emerged in the 1-4 ms lag of the raw cross-correlograms, satisfying Criterion 1 (and also Criterion 2, not shown), or a “false alarm” in the language of signal detection theory. We thus encountered two kinds of errors when utilizing exogeneous stimulation to increase neuronal firing rates – “misses” and “false alarms”, both linked to Criterion 1, due to appearance of spurious peaks in 1-4 ms lag in the cross-correlogram.

In previous studies, increase in stimulus-driven firing rate has been shown to couple with higher degree of synchronization within and across neural circuits (Temereanca et al. 2008; Wang et al. 2010) that can potentially dominate the temporal relationship of the network dynamics revealed by correlational analysis (Ginzburg and Sompolinsky 1994). This highlights the issues underlying the tradeoff between firing rate and synchronization of the local network that are intrinsic to the correlational analysis used for monosynaptic connection inference. Several paired recording studies have approached this problem by collecting spontaneous spiking data (Swadlow and Gusev 2001), where presumably the spontaneous activity is less synchronous than stimulus driven activity, or by providing weak or non-synchronizing inputs to probe the circuit (Bruno and Sakmann 2006; Bruno and Simons 2002; Reid and Alonso 1995). However, some degree of synchronization is always present, and a direct understanding of the potential effects and implications of population synchronization of presynaptic neurons on monosynaptic connection inference is lacking. To relate the amount of synchronization across the thalamic units with the increase in stimulus strength, we quantified synchrony using the spike cross-correlogram measured across thalamic pairs. Specifically, the synchrony was defined as the number of spikes of the cross-correlogram within a window of +/- 7.5 ms, normalized by the number of thalamic spikes (see Methods). Population data showed significant increases in thalamic synchrony comparing spontaneous and transient conditions, as well as sinusoidal and transient conditions (Figure 7E, n =15 pairs). The raw spike cross-correlograms for the two VPm units shown in the top row of Figure 7A are shown in Figure 7F (top and orange symbol in bottom), illustrating an increase in the central peak of the cross-correlogram and thus the thalamic synchrony, note the emergence of sharp peaks from spontaneous to transient stimulus. The nature of the stimulus thus strongly affects the synchrony, but also strongly affects the firing rate, suggesting a more general relationship between firing rate and synchrony. To more generally quantify the relationship between thalamic synchrony and mean firing rates, across several experiments we collected thalamic responses to different stimuli (spontaneous, sinusoidal (25 deg/s) and transient stimuli of different velocities (50, 125, 300, 600, 900, and 1200 deg/s), producing a range of firing rates and corresponding degrees of synchrony, displayed in Figure 7F. In general, there was a monotonic increase in thalamic synchrony with firing rate, fit well by an exponential function (Figure 7F, blue curve).

To more systematically explore the relationship between the measured thalamic synchrony and Criterion 1 of the thalamocortical connectivity inference, we generated a set of simulations based on experimental data to demonstrate how increasing degrees of thalamic synchrony could influence the connectivity inference. Described in more detail below, the simulations were based on introducing spiking activity that corresponds to varying degrees of synchrony into the experimentally observed spiking activity at the levels of both thalamus and cortex. As described above for experimental observations, through these simulations we also found that the synchronization of the presynaptic thalamic population could potentially produce errors in two scenarios: misses (i.e. a connected pair being incorrectly inferred as “not connected”, denoted C →NC) and false alarms (i.e. a not connected pair being incorrectly inferred as ‘connected’, denoted NC → C).

We first sought to simulate the scenario of the top row of Figure 7C where a pair of connected thalamocortical neurons (VPm Unit 1-S1 Unit 1) was misclassified as not-connected because a significant maximum peak was not detected in the central 1-4 ms lag of the raw cross-correlogram (violating Criterion 1, rightmost panel). We hypothesized that this experimental observation was due to a significant increase in cortical firing that was caused by inputs of nearby VPm neurons also connected to the same cortical neuron S1, but relatively synchronous with the reference thalamic neuron VPm1. For this simulation, we therefore emulated this scenario by reintroducing jittered spikes from VPm Unit 1 and S1 Unit 1 back into these datasets, respectively. Importantly, we found from our experimental observations that the mean thalamic and cortical firing rate increased with stimulus strength, but cortical firing increased to a lesser extent as compared to thalamic firing (not shown). Therefore, the number of added spikes and the jitter (zero-mean Gaussian noise of a specific standard deviation, σ) were both set to produce a specified level of synchrony between the original and perturbed VPm1 spike train and also match the observed firing rates in both VPm and S1 for this particular condition. The resultant thalamic and cortical activity was denoted as VPm Unit 1* and S1 Unit 1*, respectively. A schematic of this was shown in Figure 8A. With these manipulations, we performed the analysis of connectivity as before between VPm Unit 1* and S1 Unit 1* at each firing rate level (with corresponding thalamic synchrony level) to generate the probability of the connectivity inference (i.e. probability of satisfying Criterion 1 reported as the fraction of bootstrapped iterations results in satisfying Criterion 1). We found that the probability of inferring a connection dropped with increasing thalamic synchrony as shown in the top of Figure 8B. For low levels of synchrony (e.g. thalamic synchrony measures of 0.2 or less), the probability of satisfying Criterion 1 (and thus making the correct inference, given that Criterion 2 is satisfied) remained relatively high (from ∼0.6 to 0.9). However, increasing levels of thalamic synchrony (and certainly above 0.5), the probability of satisfying Criterion 1 approached 0 (and thus the chance of a miss neared 100%). The raw spike cross-correlogram for two synchrony levels (highlighted with the green and blue vertical dashed lines in Figure 8B top) are shown below, illustrating the disappearance of the peak in the 1-4ms lag with increased synchrony. This is consistent with what we observed in the actual experimental data (Figure 7C, top right). Note that for artificially high levels of synchrony, approaching near perfect synchrony, the probability of satisfying Criterion 1 does gradually come back up and approach the same probability as for very low levels of synchrony, as expected (not shown for simplicity).

**Figure 8:**
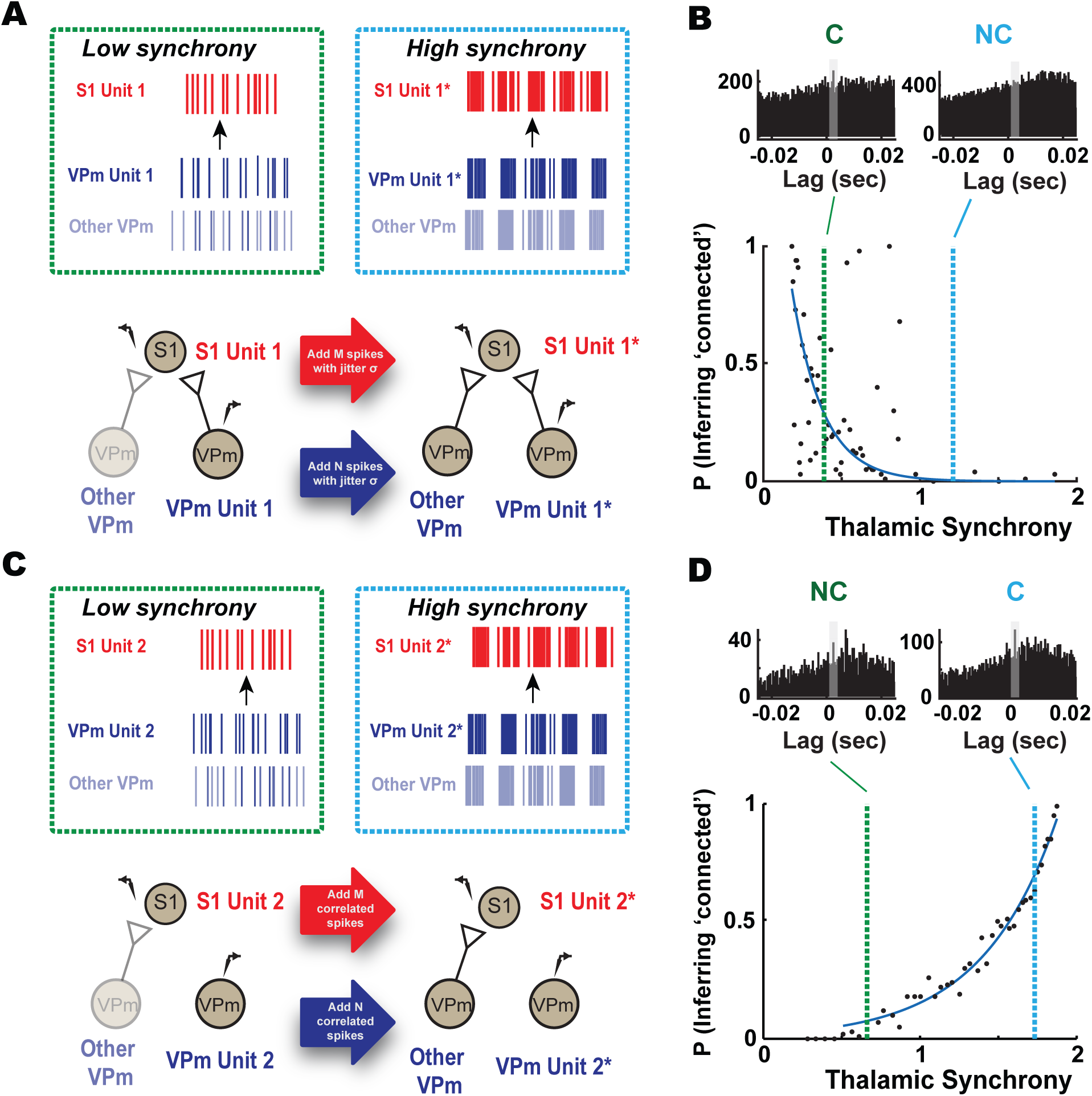
Potential errors in connectivity inference due to thalamic synchrony. **A:** Schematic shows conditions for two different synchrony levels, low and high. As thalamic synchrony increases, both thalamic and cortical showed an increase in firing rate accompanied by highly synchronous spiking in the local VPM population. To approximate this effect, we gradually added synchronous spikes in VPm and S1. We matched the firing rate of thalamic (N spikes) and cortical cells (M spikes) using experimental data and introduced the same amount of jitter (σ) associated with a specific thalamic synchrony level to both spike trains. **B.** The effects of increase in synchronous firing on the monosynaptic connectivity inference were examined with a probabilistic measure. Our simulation showed that with increasing thalamic synchrony, the probability of satisfying the criterion for monosynaptic connection decreased. Two example raw cross-correlograms, corresponding to low and high synchrony level, showed that a ‘connected’ pair (green, p = 0.3 at synchrony level of 0.25) could be misclassified as ‘not-connected’ at a higher synchrony level (light blue, p = 0.05 at synchrony level of 1.2). **C**. Same as A but for an example not connected (NC) thalamocortical pair. We gradually increased the thalamic and cortical firing by introducing synchronous spikes from a neighboring connected pair (denoted as N correlated spikes for thalamic and M correlated spikes for cortical cell). We found that the probability of error rapidly increased with thalamic synchrony (probability of satisfying Criterion 1 exceeds 0.5 as thalamic synchrony reaches 1.5. **D:** The probability of incorrectly inferring connectivity significantly increased with increasing synchrony. Two example raw cross-correlograms at the corresponding thalamic synchrony level showed that a ‘not-connected’ pair could be misclassified as ‘connected’ at a higher synchrony level. Note the emergence of ‘monosynaptic peaks’ in raw cross-correlograms with increased synchrony.

On the other hand, we found that in addition to causing the incorrect classification of a connected pair (i.e. a “miss”), the thalamic synchrony induced by increased stimulus drive could also have the opposite effect – the incorrect classification of a not-connected pair as connected (i.e. a “false-alarm). When considering a thalamocortical pair that is not synaptically connected (VPm Unit 2-S1 Unit 2), increased stimulus drive introduces the presence of spiking in the S1 neuron induced by nearby VPm neurons that serve as synaptic inputs to the S1 neuron in question. When these VPm neurons become increasingly synchronous with VPm Unit 2, this results in the presence of a peak in the raw cross-correlogram in the 1-4ms bin, satisfying Criterion 1 and thus resulting in a “false alarm”, displayed in Figure 7E.

For a demonstration of this phenomenon, from experimentally obtained data, we conducted a simple simulation designed to systematically explore this effect. Specifically, in addition to the not-connected pair in question, VPm Unit 2-S1 Unit 2, we identified a distinct, simultaneously recorded thalamocortical pair that was inferred to be connected. By introducing spikes from the connected thalamic and cortical neuron into the spike trains of VPm Unit 2 and S1 Unit 2, respectively, the degree of thalamic synchrony was systematically increased as a function of the number of spikes added, but at the same time introduced the presence of cortical spiking from the connected pair. As with the first simulation, the number of spikes added to VPm Unit 2 and S1 Unit 2 was set to match the observed thalamic and cortical firing rates for this condition, to produce the perturbed spikes trains VPm Unit 2* and S1 Unit 2*. A schematic of this was shown in Figure 8C. With these manipulations, we performed the analysis of connectivity as before, but now between VPm Unit 2* and S1 Unit 2* at each firing level (with corresponding thalamic synchrony level) to generate the probability of the connectivity inference (i.e. probability of satisfying Criterion 1 reported as the fraction of bootstrapped iterations results in satisfying Criterion 1), as with the first simulation. We found that as thalamic firing became more similar with increasing amount of mixing (reflected in an increase in thalamic synchrony), the raw cross-correlogram exhibited the increased likelihood of a peak in the 1-4ms bin. Correspondingly, the probability of error rapidly increased with thalamic synchrony (probability of satisfying Criterion 1 exceeds 0.5 as thalamic synchrony reaches 1.5 in the top panel of Figure 8D). The raw spike cross-correlogram for two synchrony levels (highlighted with the green and blue vertical dashed lines in Figure 8B right top), illustrated the appearance of the peak in the 1-4ms lag with increased synchrony (Figure 8D bottom right). Although these simulations should be considered primarily in terms of the basic trends they exhibit, the rapid increase in the probability of error with thalamic synchrony suggests that above some level of synchrony in the pre-synaptic population, false-alarms are inevitable.

## Discussion

The field of neuroscience is in a period of rapid tool development, spawned by a combination of innovative technologies and a shift in the focus of international scientific priorities. The explosion of tools for performing large scale neuronal recording at single cell resolution provides access to real-time monitoring of network activity within and across multiple brain regions (Ahrens et al. 2013; Chung et al. 2019; Jun et al. 2017). This access enables the exciting potential for interpretation of the causal flow of neuronal activity that ultimately underlies brain function, shaping of perception, and behavior (Sheikhattar et al. 2018). There have been significant efforts in anatomical tracing of connectivity and cell-type specific projections within and across brain regions and hemispheres, leading to new insights into the detailed structure of brain circuits (Yuan et al. 2015). The ultimate goal is to understand how this complex structure gives rise to behaviorally relevant function through the dynamic interaction of neurons within the neural network. However, the identification of functional connections amidst perturbation-induced confounding variables in an intact brain remains very challenging despite increasing accessibility (Lepperød et al. 2018). Here, we provided an experimental and analytical framework for quantifying long-range synaptic connectivity that was developed and tested in the thalamocortical circuit of the rodent somatosensory pathway but is generalizable to other circuits and pathways. Importantly, we attacked this from a scalable statistical framework based on signal detection theory, where we established approaches to assess confidence in classification, and systematically examined factors contributing to the inference of connectivity.

The gold standard of studying and assessing synaptic connectivity involves direct manipulation of pre-synaptic neurons to observe a measurable postsynaptic effect. Hence, connectivity studies are often performed using in-vitro brain slices (Jiang et al. 2015; Pfeffer et al. 2013) due to better accessibility to pre-and postsynaptic neurons concurrently, which is difficult or intractable in-vivo (Jouhanneau et al. 2015; Jouhanneau et al. 2018; Pala and Petersen 2015). In attempts to identify connectivity in-vivo, previous studies have used electrical stimulation to conduct collision tests that involve comparing the timing of anti-dromic activation with stimulus evoked activity to verify origins of projection (Kathleen Kelly 2001; Swadlow 1989). Although this is a powerful and attractive approach, as it helps to more confidently assess connectivity and establish causal relationships, it does not scale well to assessing connectivity at the population level, where selective stimulation of individual neurons is not typically possible. As an alternative, we adopted non-synchronizing, weak sensory drive (Bruno and Sakmann 2006; Wang et al. 2010) to assess likely connectivity through spike cross-correlation analysis. Although this approach obviously does not address issues of causality directly, it scales with increasing size of population recordings in the pre-and post-synaptic regions, opening up the possibility for assessing connectivity using large, ensemble recordings (Juavinett et al. 2018).

The systematic evaluation of the parameters of analysis presented here offers the potential for the optimization of experiment design for the purpose of assessing connectivity. Although the details likely vary across brain region and experimental preparation, the results here do suggest some general rules of thumb, perhaps the most important of which revolve around the interplay between data-length and synchrony. For example, based on our data, we estimated that about ∼9000 spikes (in terms of geometric means of total thalamic and cortical spikes) were needed to reach an inference with high certainty (95% confidence interval) in the fentanyl-anesthetized rat, consistent with recommended number of spikes used using similar stimulus conditions (Bruno and Simons 2002; Wang et al. 2010) and between 6800 and 10,000 spikes for spontaneous and sinusoidal conditions respectively in the isoflurane-anesthetized mouse (data not shown). Thus, the requirements were on the same order of magnitude for both of these cases, despite significant differences in experimental preparation (i.e. anesthesia, etc.) and differences in precise anatomical details across the species.

Synchrony across the pre-synaptic population has long been implicated as a potential confound in assessing synaptic connectivity, typically tied to false alarms (type 1 error), or the incorrect inference of connectivity in non-connected pairs (Ostojic et al. 2009). The results here supported this long-held assumption, where the artificially elevated synchrony in the VPm population resulted in the increased likelihood of satisfying Criterion 1, and importantly established ranges of measured synchrony for which this is more likely. Surprisingly, we also found that increased pre-synaptic synchrony could result in a different type of confound – a miss (type 2 error), or the incorrect inference of not-connected for connected pairs. Interestingly, this effect emerged over a relatively similar level of synchrony as determined through artificial elevation of the thalamic synchrony. As expected, spontaneous activity is typically fairly asynchronous, and in this regime, the types of errors described here are unlikely for this case. However, common drive through sensory input designed to elevate firing in otherwise sparse conditions increases the likelihood of these errors, and thus the tradeoff between increased data length and synchronization plays an important role in experimental design. This is further important in extending this approach to awake, behaving conditions, with increased firing rates, and continuous modulation in population synchrony by brain state. One alternate explanation for increased likelihood of synaptic connectivity classification errors as we moved from stimulus-free condition to highly synchronizing, transient stimuli could be due a higher rate of spike sorting errors, where elevated firing across all units (single-unit and multi-unit) could cause waveform distortion and hence cluster misclassification. Despite a slight reduction in waveform signal-to-noise ratio (4% change for VPm and 1.5% change for S1), however, we found no significant difference when we compared the waveform features of all single units such as peak-to-peak amplitudes and signal-to-noise ratios in spontaneous and stimulus-present conditions (see Supplementary Figure 6, mice data).

Although the approach here was developed with generality in mind, there are several potential limitations related to the specific details of the experimental preparation. First, the data collected in this study were from immobilized, anesthetized rodents. This enabled relatively stable and long-duration recordings of well-controlled stimulus conditions that provided insights for accurate identification of synaptic connectivity across different stimulus regimes. Although we envision the broad applicability of this approach for awake, paired recordings, future experiments are required to pinpoint the optimal experimental conditions for monosynaptic connectivity inference in awake rodents. Specifically, the relationship between mean firing rate and thalamic synchrony must be determined in this context. It was known that the baseline firing rate for both thalamic and cortical neurons are higher under wakefulness (Crochet and Petersen 2006; Urbain et al. 2015), however it is unclear if thalamic synchrony also increases monotonically with mean firing rate. Additionally, the effect of whisking is likely to confound the relationship between stimulus strength and measured synchrony explored in this study, hence confounding the monosynaptic connectivity inference. However, given the techniques and experimental parameters explored in this study, we believe that this has laid the foundation for capturing connectivity during wakefulness. Ideally, a non-synchronizing stimulus (sensory or optical) should be used to elevate the mean firing rate across the aligned thalamocortical brain regions while whisker videography should be in place to record whisker movement. Given that the effects of whisking on thalamic synchrony are unclear and could increase trial-to-trial variability, epochs of whisking should likely be excluded from the analysis for monosynaptic connectivity inference.

Second, the thalamocortical circuit is built on anatomy that is well-studied, and highly convergent, with approximately 50-100 thalamic relay neurons making synapses onto a cortical layer 4 neuron with a clear topography (Bruno and Sakmann 2006). This convergent nature of the thalamocortical projections provides significant support in assessing possible connectivity, in contrast to potential connectivity across cortical laminae or other less topographically organized projections. Third, the VPm relay neurons are reported to be uniformly excitatory, thus the pre-synaptic spiking enhances the likelihood of action potentials in the post-synaptic cortical target. Although long-range projections typically tend to be excitatory rather than inhibitory, analogous approaches should certainly be developed for inhibitory connectivity that is becoming increasingly acknowledged to play a complex and pivotal role in controlling network dynamics (Isaacson and Scanziani 2011).

With the scaling up of electrophysiological recordings comes challenges. In this work, we performed highly curated, small scale electrophysiological recordings to evaluate dynamics of VPm (potential pre-synaptic) and S1 (potential post-synaptic) neurons on a pair-by-pair basis as well as large-scale pre-and post-synaptic recordings, where any pair of neurons from the upstream and downstream structures would be a candidate for connectivity inference. What comes hand-in-hand with increasing recording yield that enables us to better answer questions about circuit function is an increasing diversity of recording quality, increasing diversity of cell type, and a wide array of possibilities for connectivity. In our hands, we found that the odds of finding putative monosynaptic connections per recording session increased by at least two-to three-fold, as illustrated in Figure 5, where 5-10 single units can be isolated from each brain region for the connectivity inference. Previous studies suggest that high signal quality can be obtained with multi-site recording due to its better detectability of extracellular feature across closely spaced contacts (Buzsáki 2004; Gray et al. 1995; Harris et al. 2000). Hence, this increase in recording yield as well as the increased probability of detecting monosynaptic connectivity with high-density probe recordings were likely due to improved single unit isolation over time, enabling reliable tracking of single units over longer recording duration (3-4 hours). Yet, in practice, handling large amounts of data with probe recordings presents challenges as most of the spike sorting algorithms to-date require manual curation to improve spike classification. This suggest that a certain amount of contamination of spikes from neighboring cells is to be expected. Overall, large-scale recording of neuronal activity will reduce the number of animals, reveal important information only available from monitoring the interaction of brain networks, but the variability inherent in these datasets drives the need for a more comprehensive approach for diverse neuronal ensembles. Traditionally, cross-correlation analysis of homogeneous, highly curated, small scale electrophysiological recordings was primarily used to attach a binary outcome for concluding monosynaptic connection in-vivo (Bruno and Simons 2002; Fujisawa et al. 2008; Miller et al. 2001; Reid and Alonso 1995; Swadlow 2003; Swadlow and Gusev 2001) and data with uncertainties were often discarded. In this work, the evaluation of the neuronal data would not result in a binary classification of connected or not, but instead the assignment of a likelihood of connectivity based on the statistical framework we present here, producing a probabilistic connectivity map. While this approach was developed through analysis of extracellular neuronal spiking data, we also envision that this would be readily adapted to optical imaging approaches that enable cellular resolution calcium or voltage imaging of population spiking activity. Recently, model-based approaches to improve connectivity estimation are being developed for the analysis of large datasets (Chen et al. 2011; Kobayashi and Kitano 2013; Kobayashi et al. 2019; Lütcke et al. 2013; Okatan et al. 2005; Paninski et al. 2004; Stevenson et al. 2008; Zaytsev et al. 2015). These methods typically involve fitting neuronal data or correlational relationships from recordings through models (i.e. generalized linear models (GLMs)) and further aim to reconstruct neuronal circuitry to estimate functional connectivity or to decode causal flow. The statistical approach we present here could eventually be combined with these more structured network modelling approaches to provide a more comprehensive framework for assessing and understanding causal interactions in the brain.

## Acknowledgements

This work was supported by NIH/NIMH Brain Initiative Grant U01MH106027 (GBS and CRF) and NIH/NINDS Brain Initiative Grant R01NS104928 (GBS). YL was supported by a Georgia Tech-Emory-PKU Global Biomedical Engineering Fellowship. AP was supported by a postdoctoral fellowships P300PA_177861 and P2ELP3_168506 from the Swiss National Science Foundation (SNSF). CJW was supported by an NIH NRSA Pre-doctoral Fellowship. WAS is supported by an NSF Graduate Research Fellowship.

---

**Supplementary Fig 1.**
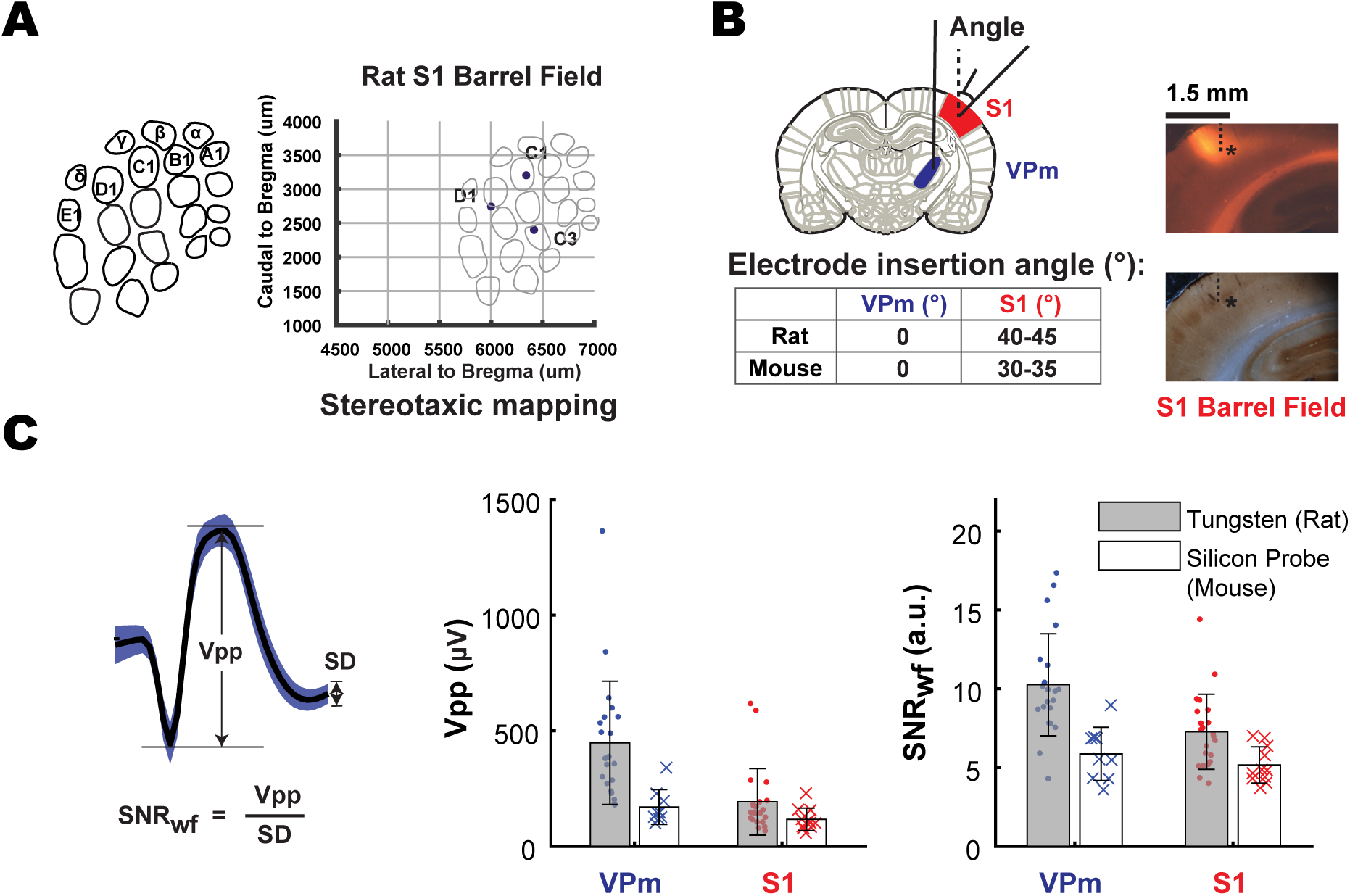
VPm-S1 targeting and unit isolation quality. **A.** S1 targeting in rat (similar concept to IOS imaging in mouse, related to main Figure 2A): Prior to actual recording, location of S1 barrels was determined using functional measure (electrophysiology) and co-registered with stereotaxic coordinates relative to bregma. We mapped three distinct locations of the barrels and fitted a barrel map template using the triangulation method. This functional map was later used to guide electrode placement for targeting a desired barrel column. **B.** The optimal electrode insertion angle for targeting VPm and S1 in rat and mouse respectively. Histological images showed fluorescent electrode track (top) targeting S1 L4 (cytochrome oxidase staining at the bottom, see Methods) in a rat. Note: * and dotted line were plotted on both images to help registering location of fluorescent signal to barrels revealed by cytochrome oxidase staining. **C.** Schematic shows the wave-form characteristics of an example single-unit. Peak-to-peak amplitude (Vpp) was defined as the voltage difference between the peak and trough of the mean waveform. Signal-to-noise ratio (SNR_wf_) of the waveform was defined as the ratio of the peak-to-peak voltage of the mean waveform divided by the standard deviation of the waveform. Single-units recorded using Tungsten microelectrodes in rats have slightly higher peak-to-peak amplitude and signal-to-noise ratio (***Vpp:*** VPm (rat): 448 ± 266 μV, n = 22 neurons; VPm (mouse): 170 ± 75.3 μV, n = 9 neurons; S1 (rat): 170 ± 75.3 μV, n = 22, S1 (mouse): 117 ± 48.6 μV, n = 11; ***SNR_wf_:*** VPm (rat): 10.3 ± 3.24 a.u., n = 22 neurons; VPm (mouse): 5.87 ± 1.69 a.u., n = 9 neurons; S1 (rat): 7.27 ± 2.38 a.u., n = 22, S1 (mouse): 5.17 ± 1.15 a.u., n = 11). All thalamic units have ISI violation <2% and cortical units have ISI violation <1%.

**Supplementary Fig 2.**
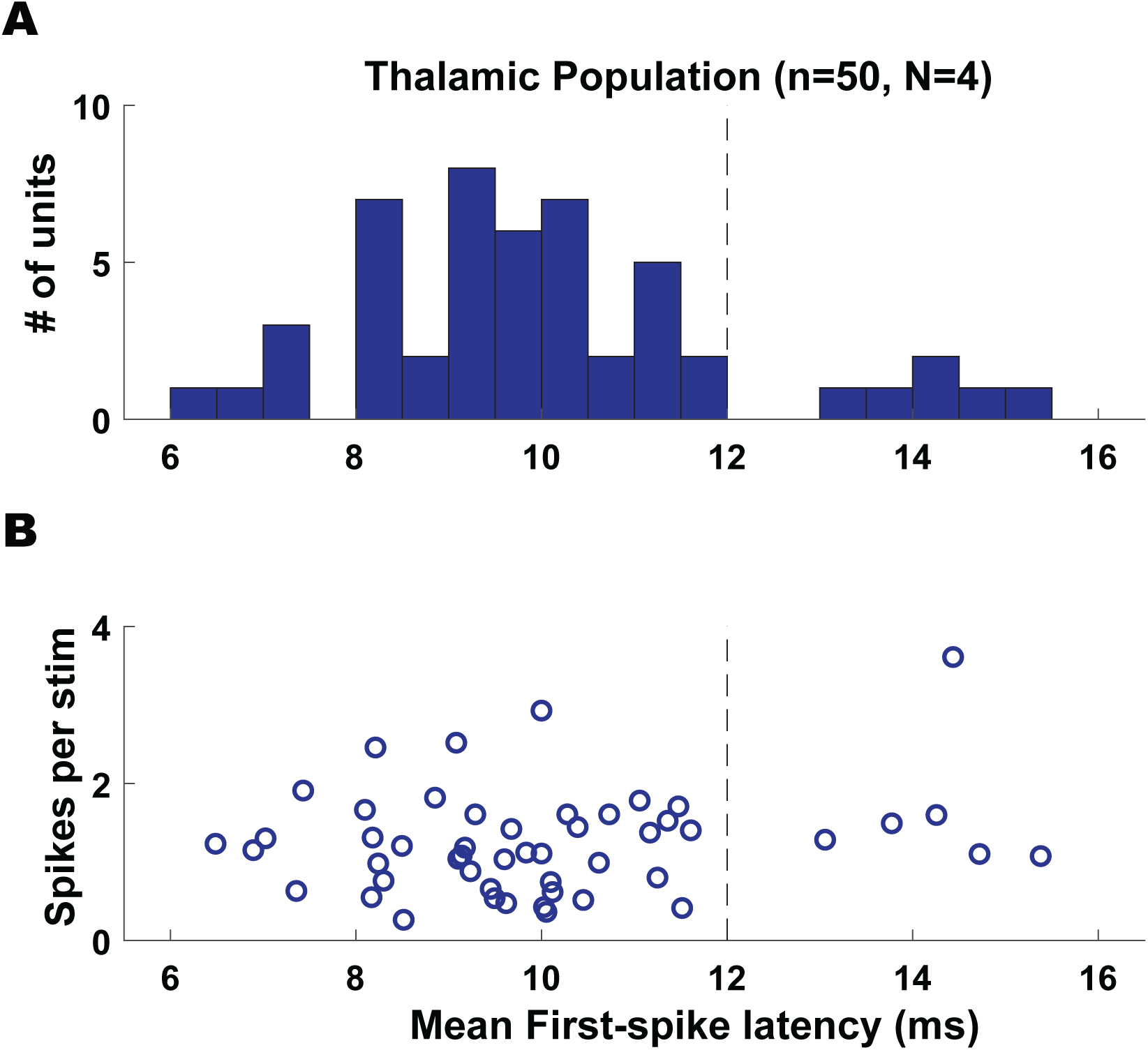
Distribution of thalamic evoked responses to punctate whisker stimulation in mice. **A.** Distribution of mean first-spike latency of evoked responses of thalamic population in response to 1st pulse of a repetitive (8-10Hz) transient stimulation (related to Figure 2D, not-adapted response). We found that majority of these cells (44 out of 50) exhibited short first-spike latency (9.43 ± 1.32 ms, mean ± SEM). However, there were a small set of cells (6 out of 50) that exhibited longer latencies (13-15 ms), which could be secondary whisker responses (more likely) or POm responses (less likely). **B.** Corresponding averaged spikecount per stimulus as a function of mean first-spike latency for all units shown in A (1.25 ± 0.66 spikes/stim, mean ± SEM, n = 50, single-unit and multi-unit data, excluding any neurons exhibiting reliability <20%). Dotted line depicted latency cut-off for VPm neuron classification (see Methods).

**Supplementary Fig 3.**
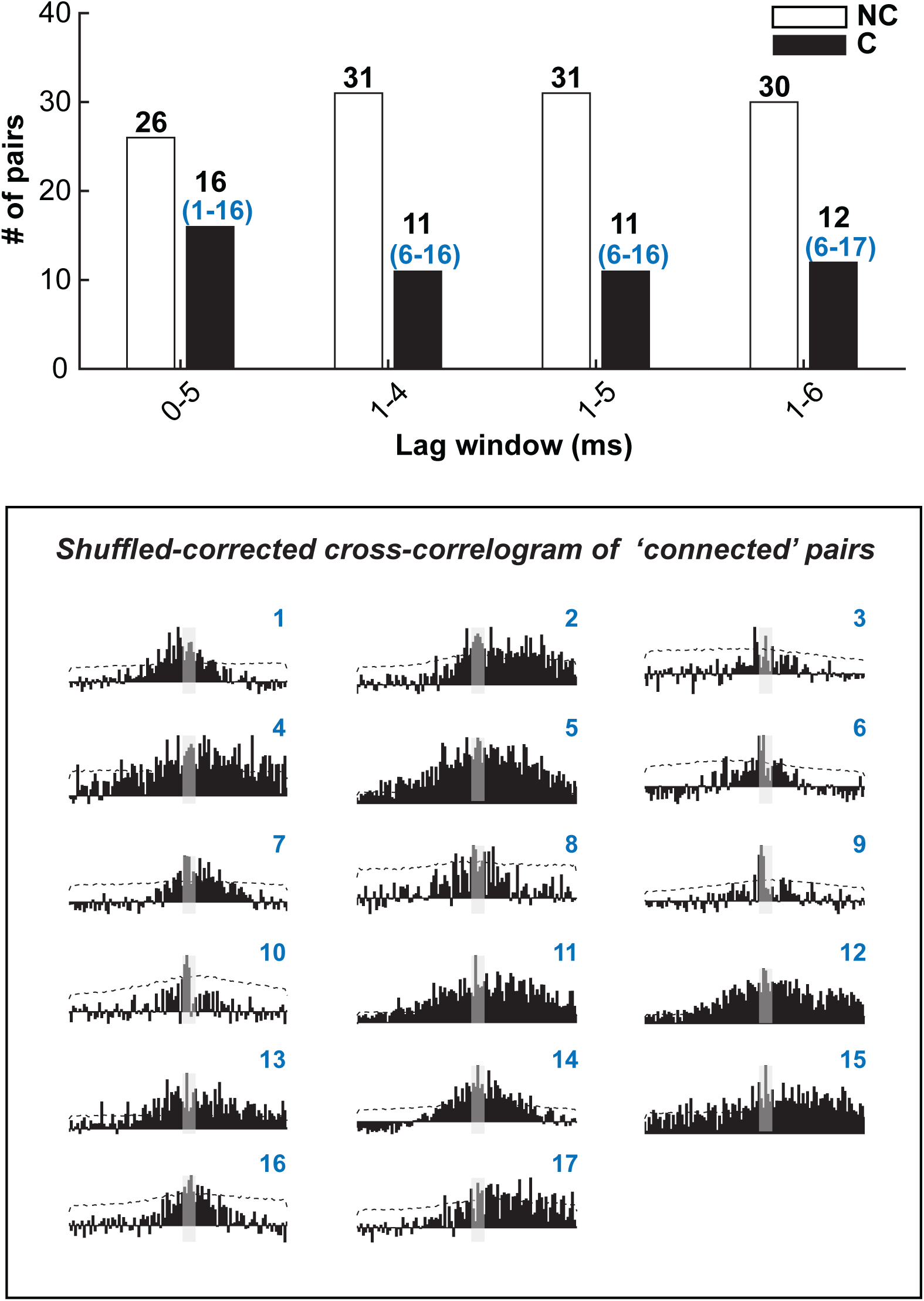
Comparison of different lag windows for connectivity analysis for rat and mouse data. **Top:** Distribution of ‘not-connected’ (NC) and ‘connected’ (C) pairs based on different selections of lag window on the cross-correlograms. The parenthesized numbers (blue) correspond to a collection of indexed cross-correlograms shown below. **Bottom:** Shuffled-corrected cross-correlograms for ‘connected’ pairs, pooled across all four lag windows. Note that each cross-correlogram is indexed from 1-17, depicting the total number of ‘connected’ pairs by any definition of lag windows. Note that cross-correlograms (index: 6-16) are the same pairs that were classified as ‘connected’ in Figure 4E, also shown in Supplementary Figure 4).

**Supplementary Fig 4.**
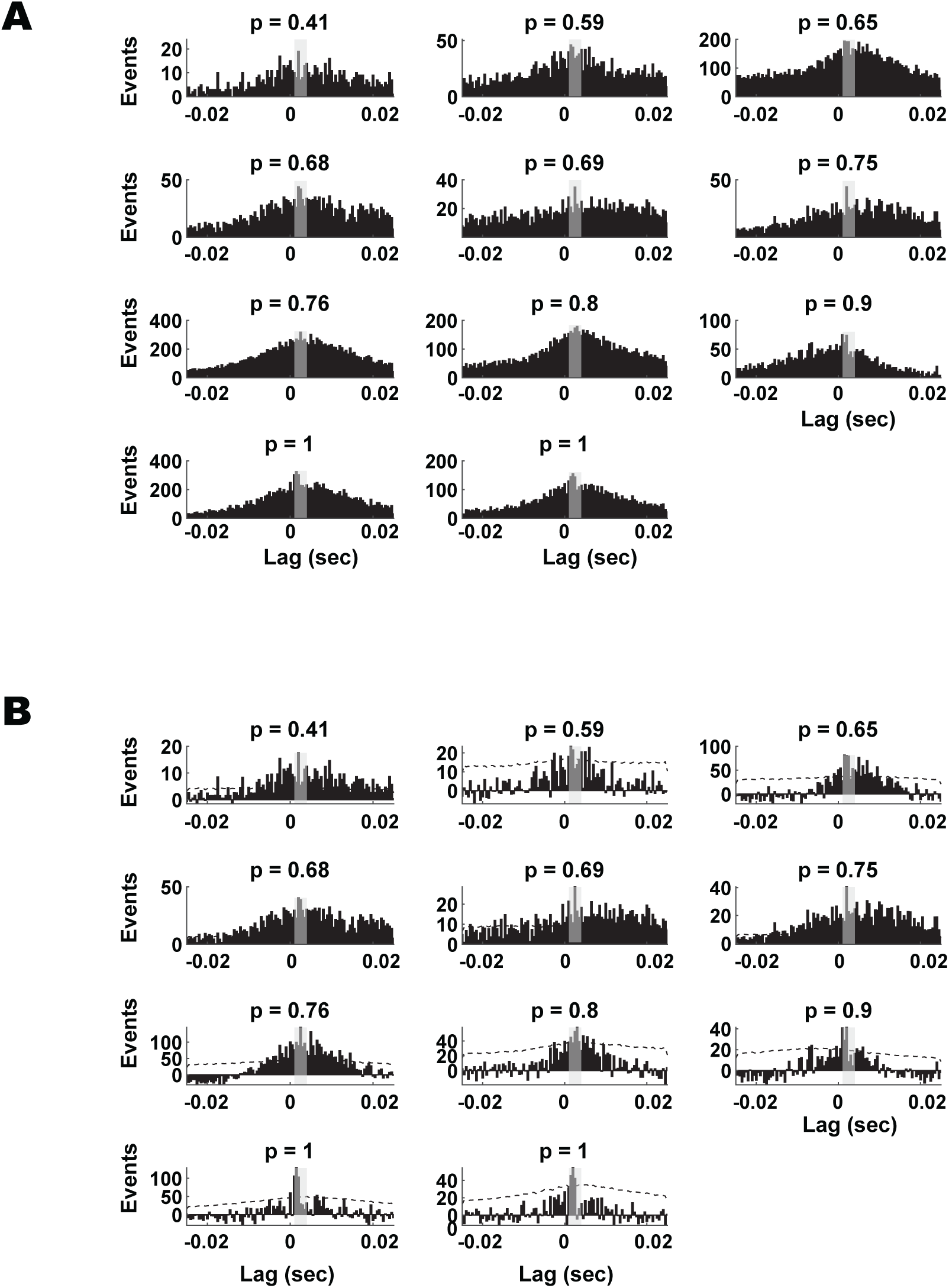
Raw and shuffled-corrected cross-correlograms of ‘connected’ pairs in rat and mouse. **A.** Raw cross-correlograms of all ‘connected’ pairs in Figure 4E. **B.** Corresponding shuffled-corrected cross-correlograms in A. P-value above each plot represents the probability of a correct inference associated with each ‘connected’ pair (related to Figure 4 and Figure 5).

**Supplementary Fig 5.**
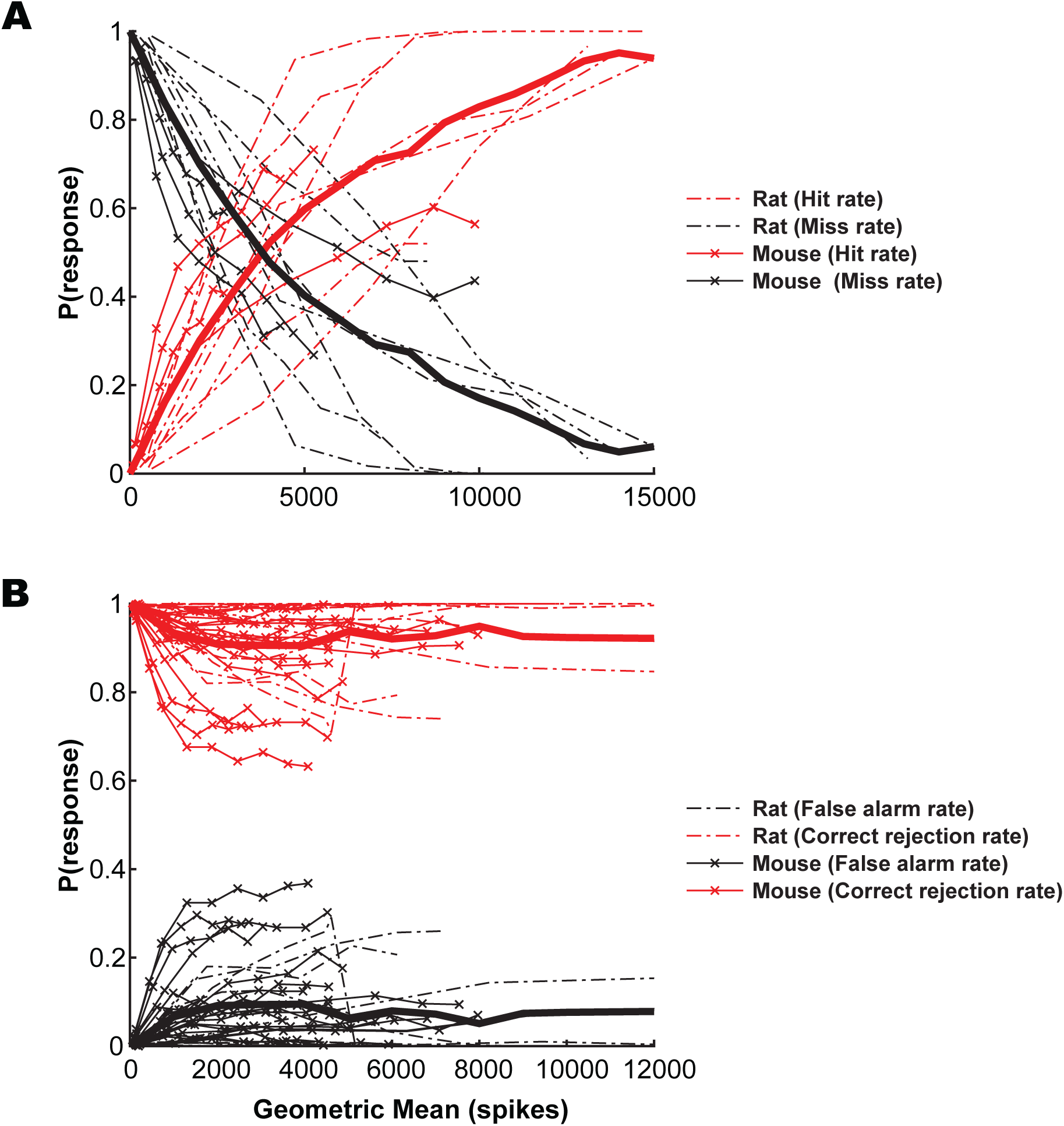
Effect of data-length on monosynaptic connectivity inference for rat and mouse data. **A.** Hit and miss rate for all connected pairs (n = 11 pairs). **B.** Correct Reject and False Alarm rate for all not-connected pairs (n = 31 pairs). Note that the hit rate and correct reject rate are similar to P(correct inference) on other figures (related to Figure 6). Bold lines represent mean probabilities, averaged across pairs at each data-length (if available).

**Supplementary Fig 6.**
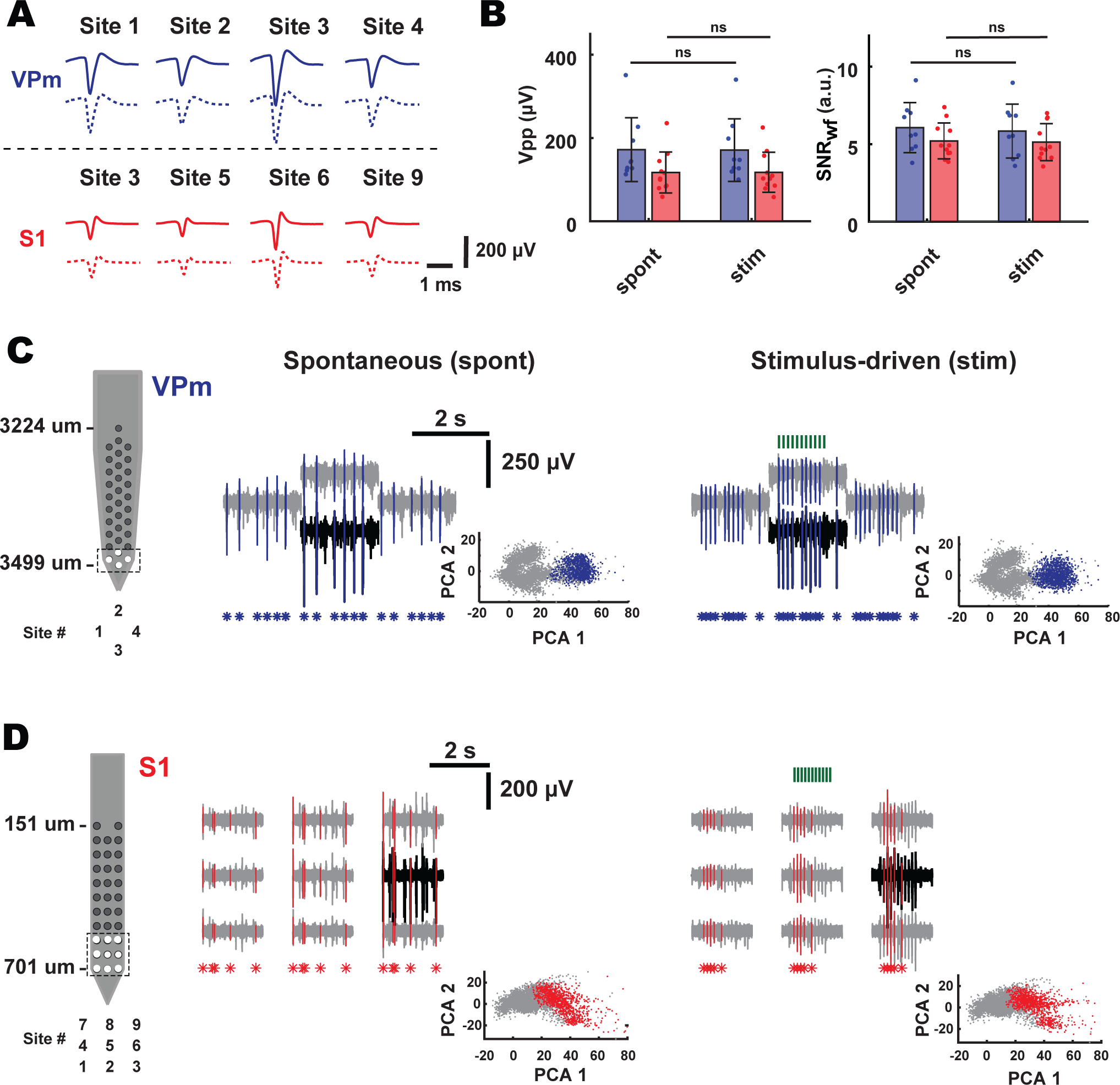
Single-unit isolation quality of probe recordings during spontaneous and stimulus-driven conditions. **A.** Example mean single-unit waveforms in C and D for spontaneous (top, solid-filled) and stimulus-driven (bottom, dashed-filled) conditions. **B.** Peak-to-peak amplitude and unit quality of waveform across spontaneous and stimulus-driven condition. (Spontaneous (Vpp): VPm = 172 ± 76.7 μV, n = 9 neurons, S1 = 117 ± 49.3 μV, n = 11 neurons; Stimulus-driven: VPm = 171 ± 75.2 μV, n = 7 neurons, S1 = 118 ± 48.2 μV, n = 11 neurons; Spontaneous (SNR_wf_): VPm = 6.06 ± 1.62 arbitrary unit (a.u.), n = 9 neurons, S1 = 5.21 ± 1.16 a.u., n = 11 neurons; Stimulus-driven: VPm = 5.84 ± 1.74 a.u., n = 9 neurons, S1 = 5.13 ± 1.19 a.u., n = 11 neurons, N = 1 mouse) **C.** Electrode configuration shown on the left. Sample voltage trace from VPm single unit with detected spikes (highlighted in blue) for spontaneous and stimulus-driven conditions. Inset (bottom): Waveform feature scatterplot showing spike clusters of the example unit and noise (gray, spikes from other whisker-responsive units detected on the same or neighboring channels). **D.** same as (C) but for S1 single unit. PCA 1 and PCA 2 plotted here are the two largest principle components of the example unit and other spikes are projected onto these dimensions.

## References

Ahissar E, Sosnik R, and Haidarliu S. Transformation from temporal to rate coding in a somatosensory thalamocortical pathway. Nature 406: 302–306, 2000.

Ahrens MB, Orger MB, Robson DN, Li JM, and Keller PJ. Whole-brain functional imaging at cellular resolution using light-sheet microscopy. Nature Methods 10: 413–420, 2013.

Alonso J-M, and Martinez LM. Functional connectivity between simple cells and complex cells in cat striate cortex. Nat Neurosci 1: 395–403, 1998.

Alonso J-M, Usrey WM, and Reid RC. Rules of Connectivity between Geniculate Cells and Simple Cells in Cat Primary Visual Cortex. The Journal of Neuroscience 21: 4002, 2001.

Barthó P, Hirase H, Monconduit L, Zugaro M, Harris KD, and Buzsáki G. Characterization of Neocortical Principal Cells and Interneurons by Network Interactions and Extracellular Features. Journal of neurophysiology 92: 600–608, 2004.

Berényi A, Somogyvári Z, Nagy AJ, Roux L, Long JD, Fujisawa S, Stark E, Leonardo A, Harris TD, and Buzsáki G. Large-scale, high-density (up to 512 channels) recording of local circuits in behaving animals. Journal of Neurophysiology 111: 1132–1149, 2014.

Borden PY, Ortiz AD, Waiblinger C, Sederberg AJ, Morrissette AE, Forest CR, Jaeger D, and Stanley GB. Genetically expressed voltage sensor ArcLight for imaging large scale cortical activity in the anesthetized and awake mouse (erratum). Neurophotonics 4: 039801, 2017.

Bruno RM, and Sakmann B. Cortex is driven by weak but synchronously active thalamocortical synapses. Science 312: 1622–1627, 2006.

Bruno RM, and Simons DJ. Feedforward mechanisms of excitatory and inhibitory cortical receptive fields. The Journal of neuroscience : the official journal of the Society for Neuroscience 22: 10966–10975, 2002.

Buzsáki G. Large-scale recording of neuronal ensembles. Nat Neurosci 7: 446–451, 2004.

Buzsáki G, Stark E, Berényi A, Khodagholy D, Kipke DR, Yoon E, and Wise KD. Tools for probing local circuits: high-density silicon probes combined with optogenetics. Neuron 86: 92–105, 2015.

Castejon C, Barros-Zulaica N, and Nuñez A. Control of somatosensory cortical processing by thalamic posterior medial nucleus: a new role of thalamus in cortical Function. PloS one 11: e0148169, 2016.

Chen Z, Putrino DF, Ghosh S, Barbieri R, and Brown EN. Statistical Inference for Assessing Functional Connectivity of Neuronal Ensembles With Sparse Spiking Data. IEEE Transactions on Neural Systems and Rehabilitation Engineering 19: 121–135, 2011.

Chung JE, Joo HR, Fan JL, Liu DF, Barnett AH, Chen S, Geaghan-Breiner C, Karlsson MP, Karlsson M, Lee KY, Liang H, Magland JF, Pebbles JA, Tooker AC, Greengard LF, Tolosa VM, and Frank LM. High-Density, Long-Lasting, and Multi-region Electrophysiological Recordings Using Polymer Electrode Arrays. Neuron 101: 21–31.e25, 2019.

Constantinople CM, and Bruno RM. Deep Cortical Layers Are Activated Directly by Thalamus. Science 340: 1591–1594, 2013.

Crochet S, and Petersen CCH. Correlating whisker behavior with membrane potential in barrel cortex of awake mice. Nat Neurosci 9: 608–610, 2006.

Csicsvari J, Hirase H, Czurko A, and Buzsáki G. Reliability and state dependence of pyramidal cell-interneuron synapses in the hippocampus: an ensemble approach in the behaving rat. Neuron 21: 179–189, 1998.

Diamond ME, Armstrong-James M, and Ebner FF. Somatic sensory responses in the rostral sector of the posterior group (POm) and in the ventral posterior medial nucleus (VPM) of the rat thalamus. Journal of Comparative Neurology 318: 462–476, 1992.

English DF, McKenzie S, Evans T, Kim K, Yoon E, and Buzsáki G. Pyramidal Cell-Interneuron Circuit Architecture and Dynamics in Hippocampal Networks. Neuron 96: 505–520.e507, 2017.

Fiáth R, Márton AL, Mátyás F, Pinke D, Márton G, Tóth K, and Ulbert I. Slow insertion of silicon probes improves the quality of acute neuronal recordings. Scientific Reports 9: 111, 2019.

Fiáth R, Raducanu BC, Musa S, Andrei A, Lopez CM, van Hoof C, Ruther P, Aarts A, Horváth D, and Ulbert I. A silicon-based neural probe with densely-packed low-impedance titanium nitride microelectrodes for ultrahigh-resolution in vivo recordings. Biosensors and Bioelectronics 106: 86–92, 2018.

Franklin K, and Paxinos G. The mouse brain in stereotaxic coordinates, compact. The coronal plates and diagrams. Amsterdam: Elsevier Academic Press, 2008.

Fujisawa S, Amarasingham A, Harrison MT, and Buzsaki G. Behavior-dependent short-term assembly dynamics in the medial prefrontal cortex. Nat Neurosci 11: 823–833, 2008.

Furuta T, Kaneko T, and Deschenes M. Septal neurons in barrel cortex derive their receptive field input from the lemniscal pathway. The Journal of neuroscience : the official journal of the Society for Neuroscience 29: 4089–4095, 2009.

Gil Z, and Amitai Y. Properties of Convergent Thalamocortical and Intracortical Synaptic Potentials in Single Neurons of Neocortex. The Journal of Neuroscience 16: 6567–6578, 1996.

Ginzburg II, and Sompolinsky H. Theory of correlations in stochastic neural networks. Physical review E, Statistical physics, plasmas, fluids, and related interdisciplinary topics 50: 3171–3191, 1994.

Gollnick CA, Millard DC, Ortiz AD, Bellamkonda RV, and Stanley GB. Response reliability observed with voltage-sensitive dye imaging of cortical layer 2/3: the probability of activation hypothesis. Journal of neurophysiology 115: 2456–2469, 2016.

Gray CM, Maldonado PE, Wilson M, and McNaughton B. Tetrodes markedly improve the reliability and yield of multiple single-unit isolation from multi-unit recordings in cat striate cortex. Journal of neuroscience methods 63: 43–54, 1995.

Harris KD, Henze DA, Csicsvari J, Hirase H, and Buzsáki G. Accuracy of tetrode spike separation as determined by simultaneous intracellular and extracellular measurements. J Neurophysiol 84: 401–414, 2000.

Isaacson JS, and Scanziani M. How inhibition shapes cortical activity. Neuron 72: 231–243, 2011.

Jiang X, Shen S, Cadwell CR, Berens P, Sinz F, Ecker AS, Patel S, and Tolias AS. Principles of connectivity among morphologically defined cell types in adult neocortex. Science 350: 2015.

Jouhanneau J-S, Kremkow J, Dorrn AL, and Poulet JFA. In Vivo Monosynaptic Excitatory Transmission between Layer 2 Cortical Pyramidal Neurons. Cell Rep 13: 2098–2106, 2015.

Jouhanneau J-S, Kremkow J, and Poulet JFA. Single synaptic inputs drive high-precision action potentials in parvalbumin expressing GABA-ergic cortical neurons in vivo. Nature Communications 9: 1540, 2018.

Jouhanneau J-S, and Poulet JFA. Multiple Two-Photon Targeted Whole-Cell Patch-Clamp Recordings From Monosynaptically Connected Neurons in vivo. Frontiers in Synaptic Neuroscience 11: 2019.

Juavinett AL, Bekheet G, and Churchland AK. Chronically-implanted Neuropixels probes enable high yield recordings in freely moving mice. bioRxiv 406074, 2018.

Jun JJ, Steinmetz NA, Siegle JH, Denman DJ, Bauza M, Barbarits B, Lee AK, Anastassiou CA, Andrei A, Aydın Ç, Barbic M, Blanche TJ, Bonin V, Couto J, Dutta B, Gratiy SL, Gutnisky DA, Häusser M, Karsh B, Ledochowitsch P, Lopez CM, Mitelut C, Musa S, Okun M, Pachitariu M, Putzeys J, Rich PD, Rossant C, Sun W-l, Svoboda K, Carandini M, Harris KD, Koch C, O’Keefe J, and Harris TD. Fully integrated silicon probes for high-density recording of neural activity. Nature 551: 232–236, 2017.

Kathleen Kelly GEC, Jed A. Hartings, Daniel J. Simons M. Axonal conduction properties of antidromically identified neurons in rat barrel cortex. Somatosensory & motor research 18: 202–210, 2001.

Kobayashi R, and Kitano K. Impact of network topology on inference of synaptic connectivity from multi-neuronal spike data simulated by a large-scale cortical network model. Journal of computational neuroscience 35: 109–124, 2013.

Kobayashi R, Kurita S, Kurth A, Kitano K, Mizuseki K, Diesmann M, Richmond BJ, and Shinomoto S. Reconstructing neuronal circuitry from parallel spike trains. Nature Communications 10: 4468, 2019.

Kodandaramaiah SB, Flores FJ, Holst GL, Singer AC, Han X, Brown EN, Boyden ES, and Forest CR. Multi-neuron intracellular recording in vivo via interacting autopatching robots. Elife 7: e24656, 2018.

Landisman CE, and Connors BW. VPM and PoM nuclei of the rat somatosensory thalamus: intrinsic neuronal properties and corticothalamic feedback. Cereb Cortex 17: 2853–2865, 2007.

Lefort S, Tomm C, Floyd Sarria JC, and Petersen CC. The excitatory neuronal network of the C2 barrel column in mouse primary somatosensory cortex. Neuron 61: 301–316, 2009.

Lepperød ME, Stöber T, Hafting T, Fyhn M, and Kording KP. Inferring causal connectivity from pairwise recordings and optogenetics. bioRxiv 463760, 2018.

Lien AD, and Scanziani M. Cortical direction selectivity emerges at convergence of thalamic synapses. Nature 558: 80–86, 2018.

London M, Roth A, Beeren L, Häusser M, and Latham PE. Sensitivity to perturbations in vivo implies high noise and suggests rate coding in cortex. Nature 466: 123–127, 2010.

Lütcke H, Gerhard F, Zenke F, Gerstner W, and Helmchen F. Inference of neuronal network spike dynamics and topology from calcium imaging data. Frontiers in Neural Circuits 7: 2013.

Mainen Z, and Sejnowski T. Reliability of spike timing in neocortical neurons. Science 268: 1503–1506, 1995.

Masri R, Bezdudnaya T, Trageser JC, and Keller A. Encoding of stimulus frequency and sensor motion in the posterior medial thalamic nucleus. J Neurophysiol 100: 681–689, 2008.

Matsumura M, Chen D-f, Sawaguchi T, Kubota K, and Fetz EE. Synaptic Interactions between Primate Precentral Cortex Neurons Revealed by Spike-Triggered Averaging of Intracellular Membrane Potentials *In Vivo*. The Journal of Neuroscience 16: 7757, 1996.

Millard DC, and Stanley GB. Anatomically based Bayesian decoding of the cortical response to intracortical microstimulation. In: 2013 6th International IEEE/EMBS Conference on Neural Engineering (NER)2013, p. 1457–1460.

Miller LM, Escabi MA, Read HL, and Schreiner CE. Functional convergence of response properties in the auditory thalamocortical system. Neuron 32: 151–160, 2001.

Nowak L, and J B. Cross correlograms for neuronal spike trains. Different types of temporal correlation in neocortex, their origin and significance. 2000, p. 53–96.

Okatan M, Wilson MA, and Brown EN. Analyzing functional connectivity using a network likelihood model of ensemble neural spiking activity. Neural computation 17: 1927–1961, 2005.

Ostojic S, Brunel N, and Hakim V. How Connectivity, Background Activity, and Synaptic Properties Shape the Cross-Correlation between Spike Trains. The Journal of Neuroscience 29: 10234–10253, 2009.

Pala A, and Petersen CC. In vivo measurement of cell-type-specific synaptic connectivity and synaptic transmission in layer 2/3 mouse barrel cortex. Neuron 85: 68–75, 2015.

Paninski L, Pillow JW, and Simoncelli EP. Maximum likelihood estimation of a stochastic integrate-and-fire neural encoding model. Neural computation 16: 2533–2561, 2004.

Paxinos G, and Watson C. The rat brain in stereotaxic coordinates. Qingchuan Zhuge translate 32: 2007.

Perkel DH, Gerstein GL, and Moore GP. Neuronal spike trains and stochastic point processes. II. Simultaneous spike trains. Biophysical journal 7: 419–440, 1967.

Pfeffer CK, Xue M, He M, Huang ZJ, and Scanziani M. Inhibition of inhibition in visual cortex: the logic of connections between molecularly distinct interneurons. Nat Neurosci 16: 1068–1076, 2013.

Raducanu BC, Yazicioglu RF, Lopez CM, Ballini M, Putzeys J, Wang S, Andrei A, Welkenhuysen M, Helleputte Nv, Musa S, Puers R, Kloosterman F, Hoof Cv, and Mitra S. Time multiplexed active neural probe with 678 parallel recording sites. In: 2016 46th European Solid-State Device Research Conference (ESSDERC)2016, p. 385–388.

Reid RC, and Alonso JM. Specificity of monosynaptic connections from thalamus to visual cortex. Nature 378: 281–284, 1995.

Rios G, Lubenov EV, Chi D, Roukes ML, and Siapas AG. Nanofabricated Neural Probes for Dense 3-D Recordings of Brain Activity. Nano Letters 16: 6857–6862, 2016.

Rossant C, Kadir SN, Goodman DFM, Schulman J, Hunter MLD, Saleem AB, Grosmark A, Belluscio M, Denfield GH, Ecker AS, Tolias AS, Solomon S, Buzsaki G, Carandini M, and Harris KD. Spike sorting for large, dense electrode arrays. Nat Neurosci 19: 634–641, 2016.

Sedigh-Sarvestani M, Vigeland L, Fernandez-Lamo I, Taylor MM, Palmer LA, and Contreras D. Intracellular, *In Vivo*, Dynamics of Thalamocortical Synapses in Visual Cortex. The Journal of Neuroscience 37: 5250, 2017.

Sheikhattar A, Miran S, Liu J, Fritz JB, Shamma SA, Kanold PO, and Babadi B. Extracting neuronal functional network dynamics via adaptive Granger causality analysis. Proceedings of the National Academy of Sciences 115: E3869, 2018.

Sitnikova EY, and Raevskii VV. The lemniscal and paralemniscal pathways of the trigeminal system in rodents are integrated at the level of the somatosensory cortex. Neuroscience and behavioral physiology 40: 325–331, 2010.

Sosnik R, Haidarliu S, and Ahissar E. Temporal frequency of whisker movement. I. Representations in brain stem and thalamus. Journal of Neurophysiology 86: 339–353, 2001.

Stevenson IH, Rebesco JM, Miller LE, and Körding KP. Inferring functional connections between neurons. Curr Opin Neurobiol 18: 582–588, 2008.

Storchi R, Bale MR, Biella GEM, and Petersen RS. Comparison of latency and rate coding for the direction of whisker deflection in the subcortical somatosensory pathway. Journal of neurophysiology 108: 1810–1821, 2012.

Stoy WA, Kolb I, Holst GL, Liew Y, Pala A, Yang B, Boyden ES, Stanley GB, and Forest CR. Robotic navigation to subcortical neural tissue for intracellular electrophysiology in vivo. Journal of Neurophysiology 118: 1141–1150, 2017.

Swadlow H, Waxman S, and Rosene D. Latency variability and the identification of antidromically activated neurons in mammalian brain. Experimental brain research 32: 439–443, 1978.

Swadlow HA. Efferent neurons and suspected interneurons in S-1 vibrissa cortex of the awake rabbit: receptive fields and axonal properties. Journal of Neurophysiology 62: 288–308, 1989.

Swadlow HA. Fast-spike interneurons and feedforward inhibition in awake sensory neocortex. Cerebral Cortex 13: 25–32, 2003.

Swadlow HA, and Gusev AG. The impact of ‘bursting’ thalamic impulses at a neocortical synapse. Nat Neurosci 4: 402–408, 2001.

Temereanca S, Brown EN, and Simons DJ. Rapid changes in thalamic firing synchrony during repetitive whisker stimulation. The Journal of neuroscience : the official journal of the Society for Neuroscience 28: 11153–11164, 2008.

Urbain N, Salin PA, Libourel P-A, Comte J-C, Gentet LJ, and Petersen CC. Whisking-related changes in neuronal firing and membrane potential dynamics in the somatosensory thalamus of awake mice. Cell Rep 13: 647–656, 2015.

Wang Q, Webber RM, and Stanley GB. Thalamic synchrony and the adaptive gating of information flow to cortex. Nat Neurosci 13: 1534–1541, 2010.

Whitmire CJ, Waiblinger C, Schwarz C, and Stanley GB. Information Coding through Adaptive Gating of Synchronized Thalamic Bursting. Cell Rep 14: 795–807, 2016.

Wright NC, Borden PY, Liew YJ, Bolus MF, Stoy WM, Forest CR, and Stanley GB. Rapid Cortical Adaptation and the Role of Thalamic Synchrony During Wakefulness. bioRxiv 2020.2010.2008.331660, 2021.

Yu J, and Ferster D. Functional Coupling from Simple to Complex Cells in the Visually Driven Cortical Circuit. The Journal of Neuroscience 33: 18855, 2013.

Yuan J, Gong H, Li A, Li X, Chen S, Zeng S, and Luo Q. Visible rodent brain-wide networks at single-neuron resolution. Frontiers in Neuroanatomy 9: 2015.

Zaytsev YV, Morrison A, and Deger M. Reconstruction of recurrent synaptic connectivity of thousands of neurons from simulated spiking activity. Journal of computational neuroscience 39: 77–103, 2015.

Zhang W, and Bruno RM. High-order thalamic inputs to primary somatosensory cortex are stronger and longer lasting than cortical inputs. Elife 8: e44158, 2019.

